# Non-coding germline *GATA3* variants alter chromatin topology and contribute to pathogenesis of acute lymphoblastic leukemia

**DOI:** 10.1101/2020.02.23.961672

**Authors:** Hongbo Yang, Hui Zhang, Yu Luan, Tingting Liu, Kathryn G Roberts, Mao-xiang Qian, Bo Zhang, Wenjian Yang, Virginia Perez-Andreu, Jie Xu, Sriranga Iyyanki, Da Kuang, Shalini C. Reshmi, Julie Gastier-Foster, Colton Smith, Ching-Hon Pui, William E Evans, Stephen P Hunger, Leonidas C. Platanias, Mary V Relling, Charles G Mullighan, Mignon L Loh, Feng Yue, Jun J Yang

## Abstract

Inherited non-coding genetic variants confer significant disease susceptibility in many cancers. However, the molecular processes of by which germline variants contribute to somatic lesions are poorly understood. We performed targeted sequencing in 5,008 patients and identified a key regulatory germline variant in *GATA3* strongly associated with Philadelphia chromosome-like acute lymphoblastic leukemia (Ph-like ALL). By creating an isogenic cellular model with CRISPR-Cas9 system, we showed that this variant activated a strong enhancer that significantly upregulated *GATA3* transcription, which in turn reshaped the global chromatin accessibility and 3D genome organization. Remarkably, this genotype switch induced a chromatin loop between the *CRLF2* oncogene and a distal enhancer, similar to the somatically acquired super-enhancer hijacking event in patients. *GATA3* genotype-related alterations in transcriptional control and 3D chromatin organization were further validated in Ph-like ALL patients. Finally, we showed that *GATA3* directly regulates *CRLF2* and potentiates the oncogenic effects of JAK-STAT signaling in leukemogenesis. Altogether, our results provide evidence for a novel mechanism by which a germline non-coding variant contributes to oncogene activation epigenetic regulation and 3D genome reprogramming.

## Introduction

Acute lymphoblastic leukemia (ALL) is the most common cancer in children and there is growing evidence of inherited susceptibility to this hematological malignancy(Hunger and Mullighan, 2015; Moriyama et al., 2015; Pui et al., 2015). In particular, genome-wide association studies (GWAS) have identified at least 9 genomic loci (i.e., *CDKN2A/2B*, *IKZF1*, *ARID5B*, *CEBPE*, *PIP4K2A*-*BMI1*, *GATA3*, *TP63, LHPP, and ELK3*) with common variants that influence ALL risk(Papaemmanuil et al., 2009; Perez-Andreu et al., 2013; Sherborne et al., 2010; Trevino et al., 2009; Xu et al., 2013; Xu et al., 2015). These variants cumulatively confer a substantial increase of ALL risk(Xu et al., 2015), and explain a large proportion of the estimated heritability of this leukemia(Enciso-Mora et al., 2012). Interestingly, some ALL germline risk variants also co-segregate with specific acquired genomic abnormalities in leukemia(Perez-Andreu et al., 2013; Trevino et al., 2009; Walsh et al., 2013), suggesting intricate interactions between somatic and germline mutations during leukemogenesis. In particular, we have previously reported germline intronic variants in the *GATA3* gene associated with the risk of developing Philadelphia chromosome (Ph)-like ALL(Perez-Andreu et al., 2013), a subtype characterized by a leukemia gene expression profile resembling that of Ph-positive ALL with *BCR-ABL1* fusion(Den Boer et al., 2009; Roberts et al., 2014). Each copy of the *GATA3* risk allele increased the risk of Ph-like ALL by 3.25-fold. Because Ph-like ALL is associated with distinctive genomic lesions in the cytokine signaling pathway genes (e.g., 50% of cases harbor *CRLF2*-rearrangements)(Roberts et al., 2014), it raises the question whether germline genetic variation in *GATA3* is directly or indirectly involved in the deregulation of this pathway in ALL. So far, the exact variant that determines GWAS signal at the *GATA3* locus remains unknown and the exact molecular process by which the variant(s) contribute to Ph-like ALL pathogenesis is also unclear.

Even though GWAS have identified a plethora of variants associated with diverse human traits and diseases with varying degree of effects(MacArthur et al., 2017), there is still an extreme paucity of examples that clearly demonstrate the molecular mechanisms linking risk alleles to disease pathogenesis. The main challenge is that the majority (>90%) of the disease or trait-associated variants are located in non-coding (intronic and/or intergenic) regions of the genome whose function remains largely uncharacterized. A recent work systematically analyzed ENCODE(ENCODE-Project-Consortium, 2012) and Epigenome Roadmap(Bernstein et al., 2010) data and showed that the majority of the non-coding variants are located inside regulatory elements (e.g., promoter, enhancer, and silencer)(Maurano et al., 2012), raising the possibility that genetic variants at these sites may play a regulatory role and modulate local and/or distal gene transcription. Another challenge in dissecting the regulatory roles of the non-coding elements is how to identify their target genes, as it has been shown that enhancers can function from either upstream or downstream of their target genes, from as far as 1 million base pairs away through chromatin looping(Lettice et al., 2003). Recent high-throughput methods based on chromatin conformation capture such as Hi-C presented an unprecedented opportunity to study the effects of the non-coding elements on higher-order chromatin structure in a genome-wide fashion(Dixon et al., 2012; Lieberman-Aiden et al., 2009).

In this work, we sought to systematically identify *GATA3* variants in Ph-like ALL by targeted sequencing in 5,008 ALL patients and functionally investigate the underlying mechanism of how they affect chromatin 3-dimensional structure, influence cell signaling, and contribute to leukemogenesis.

## Results

### Identification of functional regulatory variants in *GATA3 l*oci in Ph-like ALL

To comprehensively identify ALL risk variants at the *GATA3* locus, we performed targeted sequencing of a ∼27 Kb genomic region at 10q14, encompassing exons, introns, and upstream/ downstream flanking regions of *GATA3*, in 5,008 children with ALL (including 985 patients with Ph-like ALL status ascertained, **Table S1**, **Figure S1**). A total of 1,048 variants were identified, of which 127 variants had a minor allele frequency >1% and were included in subsequent analyses (**Figure S1**). Comparing the frequency of each variant in Ph-like ALL (N=141) vs. non-Ph-like ALL (N=844), we identified three variants that were significantly associated with susceptibility to Ph-like ALL after correcting for multiple testings (*P*<1×10^-5^), all of which are non-coding. Variant rs3824662 in intron 3 showed the strongest association (*P*=1.2×10^-8^, **Figure 1A**), and multivariate analysis conditioning on this SNP revealed no independent signals (**Figure S2**). Examining the chromatin state annotations of this genomic region across 42 cell and tissue types from the Roadmap Epigenomics Project(Zhou et al., 2015), we observed that rs3824662 is aligned with a putative enhancer in the hematopoietic tissues (i.e., enrichment of H3K27ac and H3K4me1 marks with an under-representation of H3K27me3 mark, **Figures 1B and S3A**). Taken together, these results pointed to rs3824662 as the likely functional and causal variants within an enhancer element that drives the association with Ph-like ALL at the *GATA3* locus.

**Figure 1.**
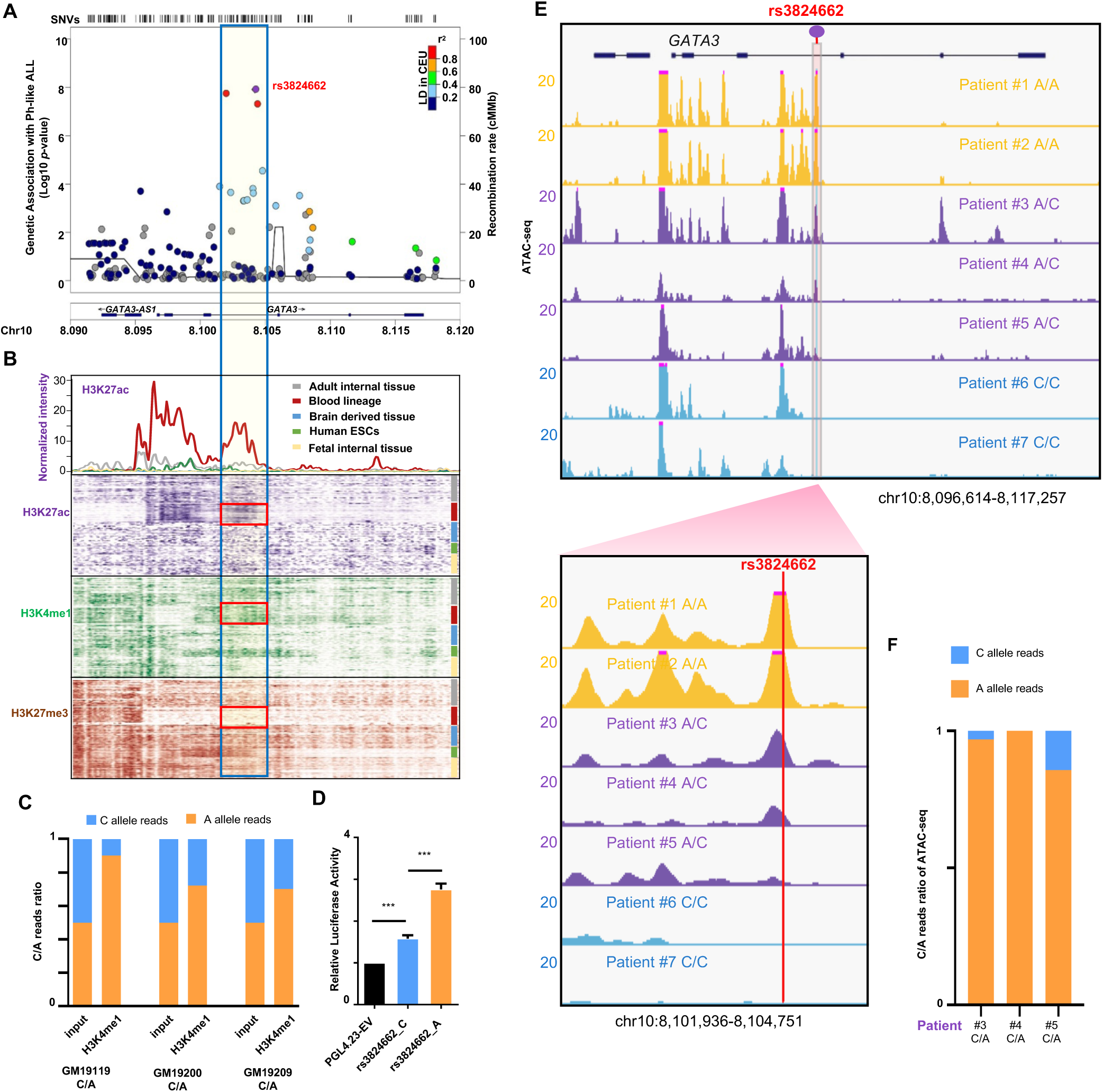
rs3824662 is the top *GATA3* variant associated with Ph-like ALL susceptibility and the leukemia risk allele (A) is associated with enhancer activity and open chromatin status. **A**, Genetic variants associated with Ph-like ALL at the *GATA3* locus discovered by targeted-sequencing. The purple dot indicates the variant with strongest association signal (rs3824662), and the blue box represents the 2.7 kb region flanking this variant. **B**, Chromatin state annotations from the Roadmap Epigenomics Project. Enhancer marks (H3K4me1 and H3K27ac) and repressive mark (H3K27me3) are plotted across 42 cell and tissue types for the *GATA3* genomic region. These epigenomic data suggest that rs3824662 is located inside a hematopoietic cell-specific enhancer element (red boxes). The upper panel shows the H3K27ac profiling in this region. The H3K27ac signals are averaged by different tissue-types and plotted in 100bp bins. **C**, Luciferase reporter assay comparing the enhancer activities of the fragments containing either the rs3824662 risk A allele or wildtype C allele in GM12878, an immortalized B lymphoblastoid cell line with normal karyotype. T bars indicate standard deviations. **D**, Allelic analysis of H3K4me1 ChIP-seq data in three lymphoblastoid cell lines with heterozygous genotype at rs3824662 (GM12119, GM19200, and GM19219). Orange and blue bars indicate the percentage of ChIP-seq reads from the A allele and the C allele, respectively. **E**, Open chromatin status at the rs3824662 locus in 7 ALL PDX samples of different genotypes, as determined using ATAC-seq. The bottom panel represents a 2.8 kb region flanking rs3824662. **F**, Allelic analysis of ATAC-seq data in three Ph-like ALL samples with heterozygous genotype at rs3824662. Orange and blue bars indicate the percentage of ATAC-seq reads from the A allele and the C allele, respectively (P3: 31 A *vs* 1 C reads, P4: 8 A *vs* 0 C reads, P5 : 6 A *vs* 1 C reads).

To validate the enhancer function of this regulatory DNA element and investigate how its activity is influenced by rs3824662 genotype, we first tested the 1,120-bp fragment surrounding rs3824662 using a reporter gene assay in lymphoblastoid cells GM12878. The wildtype fragment (with the C allele) showed a modest enhancer effect, while the same fragment with the risk A allele robustly activated reporter gene transcription with three-fold increase over the vector control (**Figure 1C**, and similar results in other cell lines shown in **Figure S3B**), suggesting that the A allele is a gain-of-function variant. Similarly, in lymphoblastoid cell lines with heterozygous genotype at rs3824662 (i.e., GM19119, GM19200, GM19209(McVicker et al., 2013)), we also observed a significant allele-biased histone modification, linking the A allele with an over-representation of the enhancer-associated H3K4me1 chromatin mark (**Figure 1D**). We then performed ATAC-Seq to profile open chromatin regions in seven primary leukemia samples from patient-derived xenografts of ALL with different rs3824662 genotypes (N=2, 3, and 2 for cases with A/A, A/C, and C/C genotype, respectively, **Table S3**). We observed that samples with the A/A genotype showed higher levels of open chromatin signals than those with A/C or C/C genotypes (**Figure 1E**). Furthermore, in three patients with heterozygous genotype at rs3824662, open chromatin signal at this locus exhibited clear allelic imbalance with the A allele preferentially linked to more chromatin accessibility (**Figure 1F**). Similarly, in a panel of B-ALL cell lines of diverse molecular subtypes, we observed that samples with the A/A genotype showed higher levels of open chromatin signals than those with C/C genotype **(Figure S3C)**. In fact, the strongest ATAC-seq signals at this locus were observed in two Ph-like ALL cell lines (MHH-CALL4 and MUTZ5), both of which have A/A genotype at rs3824662, again suggesting that the A allele was associated a more transcriptionally active chromatin state.

### rs3824662 risk A-allele upregulates *GATA3* expression

To directly assess the effects of the rs3824662 genotype, we specifically knocked in the A allele at rs3824662 in the wildtype lymphoblastoid cell line GM12878, using CRISPR-Cas9 genome editing (**Figure S4A-S4C**). Engineered GM12878 cells with the variant allele (A/C or A/A genotype) showed 3.7- and 3.8-fold increase of *GATA3* expression compared with isogenic cells with the wildtype C/C genotype (**Figure S4D**). We then performed RNA-Seq and qPCR experiments to determine whether this variant can influence gene transcription in *cis*, and we focused on genes located within the same topologically associated domains (TADs) because it has been shown that effects of cis-regulatory elements are usually confined by the TAD boundaries(Dixon et al., 2012; Hnisz et al., 2016). Of the four genes within the rs3824662-containing TAD, only the expression of *GATA3* was significantly altered upon genome editing (**Figure 2A**), further indicating that this variant specifically reguates *GATA3* transcription. Further, by analyzing the RNA-seq results of the engineered GM12878 cells with heterozygous genotype at rs3824662, we noted significant allele-biased transcription of the *GATA3* (in favor of the T allele at coding variant rs2229359 *in cis* with the A allele at rs3824662, **Figure S5A-S5C**). This allelic expression pattern confirmed the cis regulatorary effects of the rs3824662-containing enhancer. Further, we performed RNA-Seq in seven primary leukemia samples from ALL PDX and again confirmed that patients with A allele at rs3824662 is associated with higher *GATA3* expression (**Figure 2A** bottom panel). To define the target gene for this regulatory variant, we also performed Capture-C experiment to directly identify the regions that interact with this enhancer and observed that it forms a strong chromatin loop with the *GATA3* promoter (**Figure 2B**, vertical yellow bar indicates the enhancer at the rs3824662 locus and pink bar indicates *GATA3* promoter). To pinpoint the transcription factor that preferentially binds to rs3824662 risk A allele, we performed footprint analysis using the high-depth ATAC-seq data from the MHH-CALL4 cells (rs3824662 A/A allele), and identified the NFIC motif proximal to the variant (chr10: 8,104,196-8,104,208, **Figure S6A**). ChIP-qPCR of NFIC in GM12878 (WT) and GM12878 (A/A) cells also confirmed that this transcription factor preferentially bound to the A allele, at a level of 15-fold higher compared with the C allele (**Figure S6B**).

**Figure 2.**
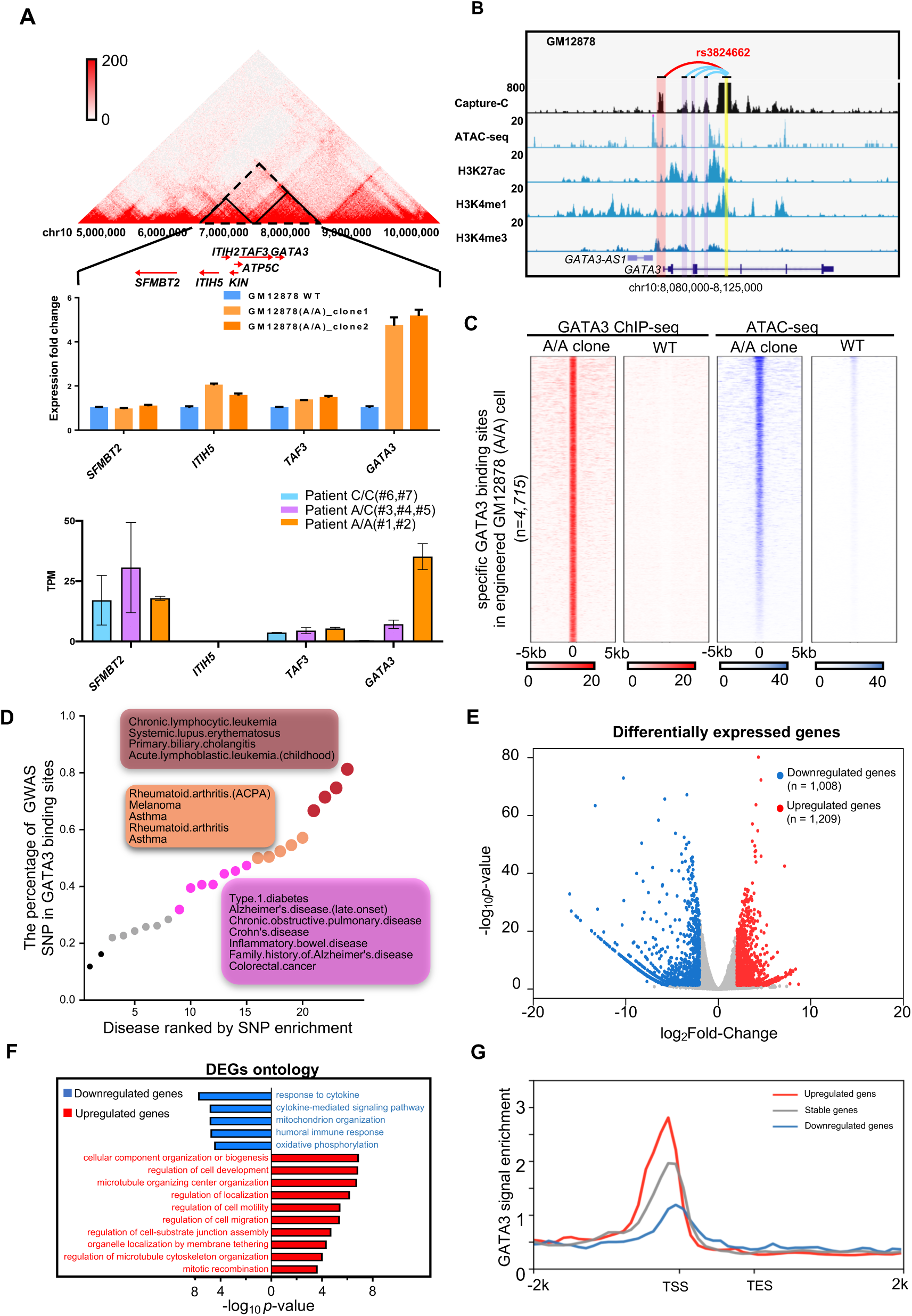
The rs3824662 A allele increases *GATA3* expression and induces global gene expression changes in GM12878 cells and ALL PDX samples. **A**, *Cis* effects of rs3824662 on gene expression within the local topologically associating domain (TAD). Gene expression was quantified by qPCR for each gene in wildtype (C/C) and engineered GM12878 cells (A/A). TAD was defined using the GM12878 wildtype Hi-C data. Expressions are normalized against *BACTIN*. Only genes that are expressed in both GM12787 lines are presented. Bottom panel shows expression of the same panel of genes (as TPM) in 7 ALL PDX samples. **B**, Chromatin interactions between the rs3824662 (yellow bar) and *GATA3* promoter (red bar) as determined by Capture-C. **C**, Heatmap of GATA3 ChIP-seq (GATA3 binding) and ATAC-seq (open chromatin status) in GM12878 (WT) and engineered GM12878 (A/A) cells. Each row represents a 6kb genomic region flanking a *GATA3* binding site that is specific in engineered GM12878 (A/A) cells. **D**, Enrichment of disease risk loci (i.e., disease GWAS hits) in GATA3 binding sites gained in engineered GM12878 (A/A) cells. **E**, Differentially expressed genes (DEGs) between GM12878 (A/A) and GM12878 (WT) cell lines (1,209 upregulated genes *vs* 1,008 downregulated genes. **F**, Gene Ontology analysis of DEGs. **G**, Genes up-regulated in the GM12878 (A/A) cell line are more significantly enriched for GATA3 binding than those not affected or up-regulated in the GM12878 (WT) cells.

### rs3824662 risk A-allele induces novel GATA3 binding sites and reshapes global chromatin accessibility landscape

Having established that the rs3824662 risk allele upregulates *GATA3* gene expression, we next sought to determine the effects of increase in GATA3 on global gene transcription and chromatin organization. Comparing genome-wide GATA3 ChIP-Seq in engineeered GM12878 cells (genotype C/C vs. A/A), we found that there was an overall increase in GATA3 binding, with 4,715 novel binding sites in the engineered A/A clones compared to isogenic cells with wildtype C/C genotype (**Figure 2C**). These GATA3 binding sites co-localized with regions that became accessible in GM12878 (A/A) cells as determined by ATAC-seq (**Figure 2C**): of the 4,715 gained GATA3 binding sites, 2,650 were also identified as novel open chromatin regions created by the A allele in GM12878. In fact, these new GATA3 binding sites were devoid of nucleosomes (**Figure S7A**), consistent with the notion that GATA3 functions as a pioneer factor(Takaku et al., 2016) and may be driving the open chromatin status at these loci. Strikingly, these novel GATA binding sites were also more likely to locate close to important Ph-like ALL genes, whose expression most strongly distinguished Ph-like ALL from other ALL subtypes(Harvey et al., 2010) (**Figure S7B**, *p*-value=0.0003, Wilcoxon test and **Figure S8A, S8B**). Interestingly, these novel GATA3 binding sites are significantly enriched for a panel of GWAS variants associated with different diseases. For example, 13 out 16 CLL-associated and 8 out of 12 ALL-associated variants are located in these novel GATA3 binding sites (**Figure 2D**). Furthermore, in the engineered GM12878 (A/A) cells, GATA3 bound to genomic loci frequently targeted by chromosomal translocations in Ph-like ALL(Roberts et al., 2014) (e.g., *CSF1R, PDGFRB*) (**Figure S9**). Globally, there are 2,217 genes are differentially expressed in the GM12878 (A/A) cell line, with 1,209 upregulated and 1,008 downregulated genes. GO term analysis showed that genes in the migration related pathways are preferentially activated in GM12878 cell line with the A/A genotype (**Figure 2F**). GATA3 binding is also signicant higher in upregulated genes, compared to downregulated genes in GM12878 (A/A) cells (**Figure 2G**, p-value<2.2e-16, Kolmogorov-Smirnov test).

### Up-regulated GATA3 leads to changes in 3D genome organization

Recent analyses using Hi-C data identified two types of compartments in the human genome with distinctive patterns of chromatin interactions: compartment A (active) and compartment B (repressive) (Dixon et al., 2012; Lieberman-Aiden et al., 2009), and the A-to-B compartment switching is associated with extensive gene expression changes. Given the role of GATA3 as a pioneer factor, we postulated that elevated *GATA3* expression (as a result of the rs3824662) would also influence 3D chromatin organization on a genome-wide scale. Therefore, we performed Hi-C experiments in GM12878 (WT) and also the engineered isogenic GM12878 (A/A) cells, and found that 4.07% of the genome underwent B-to-A compartment switch when the C allele at rs3824662 was replaced with the A allele (**Figure 3A**). Globally, B-to-A compartment switching resulted in upregulation of genes located in these regions (**Figure 3B**). Particularly notable was the *PON2* gene, which is among the most differentially expressed genes between Ph-like vs non-Ph-like ALL(Harvey et al., 2010). The *PON2* genomic locus underwent dramatic B-to-A compartment switching with a 6.258-fold increase in its expression (**Figure 3C**, upper panel), following the C-to-A allele substitution at rs3824662 in the GM12878 cell line. To further examine the functional consequences of the A allele in human primary leukemia cells, we performed Hi-C experiments in seven ALL PDX samples with different rs3824662 genotypes. Similar to what we observed in the GM12878 cell lines, we found that B-to-A compartment switch at the *PON2* locus was prominent in leukemia samples with the A/A genotype, along with transcription activation of the *PON2* gene (**Figure 3C**, bottom panel), whereas this region appeared transcriptionally inactive in WT patients. Leukemia cells with heterozygous genotype at rs3824662 exhibited intermediate phenotypes in this regard. Interestingly, Patient #4 who has a heterozygous genotype at rs3824662 showed a dramatic A compartment expansion, likely due to acquired translocation events in chr7 (**Figure S10A and S10B**), pointing to chromatin reorganization arising from somatic genomic abnormalities. Finally, ALL PDX samples containing the A allele clusterd together based on whole-genome A/B compartment states (**Figure 3D**).

**Figure 3.**
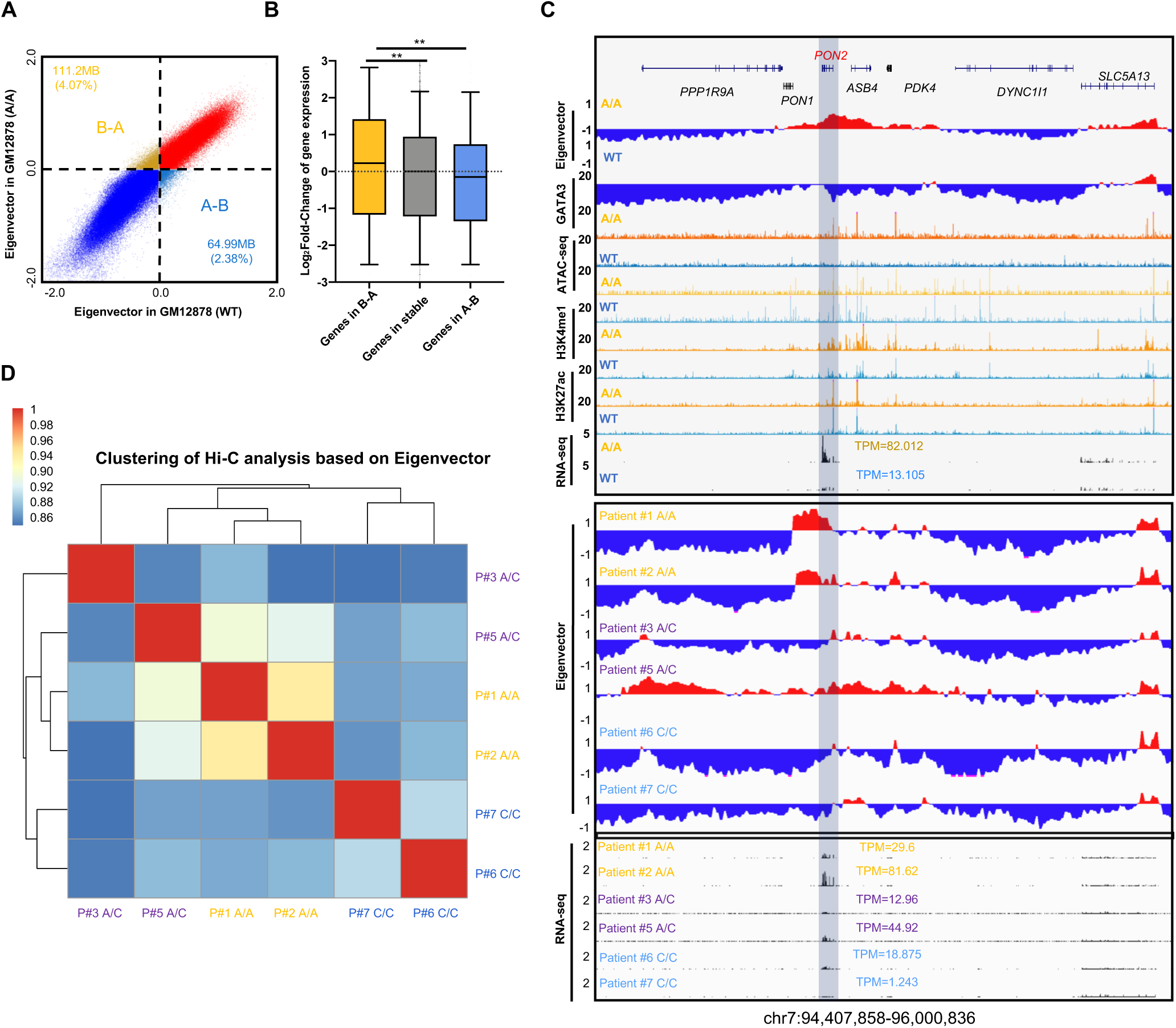
Upregulation of *GATA3* expression leads to genome-wide A-B compartment reorganization. **A**, Engineered GM12878 (A/A) cells contain more active domains (Compartment A) than GM12878 (WT) cells (1,192,100,000 bp *vs* 1,145,890,000 bp). **B**, Genes located within regions that underwent the B-to-A compartment switch showed increased expression (wildtype vs A/A genotype, *p* value < 2.2e-16 by Wilcoxon test). **C**, Ph-like ALL associated gene *PON2* locus underwent B-to-A switch in the engineered GM12878 (A/A) cells, with a 6.258-fold increase in *PON2* expression (upper panel). ALL PDX samples with risk A alleles also shows similar B-to-A switch in PON2 locus (bottom panel). **D**, Genome-wide pattern of A/B compartment states in ALL PDX samples clustered according to genotype at rs3824662 (Pearson correlation coefficient). Pearson Correlation Coefficient matrix was generated based on the A/B compartment states using 10kb resolution. A compartments were defined as 1, and B compartments were defined as −1. Grey bar indicates *PON2* gene

Although there was no signicant genome-wide change at TAD level (**Figure S11A and S11B**), we observed a set of chromatin loops in engineered GM12878 cells (A/A allele) and these loops are significantly enriciched for GATA3 binding sites (**Figure 4A**). These novel interactions in GM12878 (A/A) cells also have longer interaction distance and are enriched with higher enhancer-promoter and promoter-promoter interaction, compared to GM12878 (WT) cells (**Figure 4B, 4C and S12**). Next, we examined the chromatin interactions for the *CRLF2* oncogene and found they formed a new loop that brought the *CRLF2* promoter to close proximity to a distal super enhancer in *P2RY8* with concomitant GATA3 binding (**Figure 4D**), which may have contributed to the increase of *CRLF2* transcription in the engineered A allele cells. This new interaction between *P2RY8* and *CRLF2* is also specifically detected in ALL patient PDX samples with risk-A alleles (**Figure 4E**). Strikingly, this new linkage between the CRLF2 promoter and distal enhancer echos an enhancer hijacking event induced by an intrachromosomal rearrangment, which is one of the main mechanisms of *CRLF2* overexpression observed in ∼25% cases of Ph-like ALL(Roberts et al., 2014).

**Figure 4.**
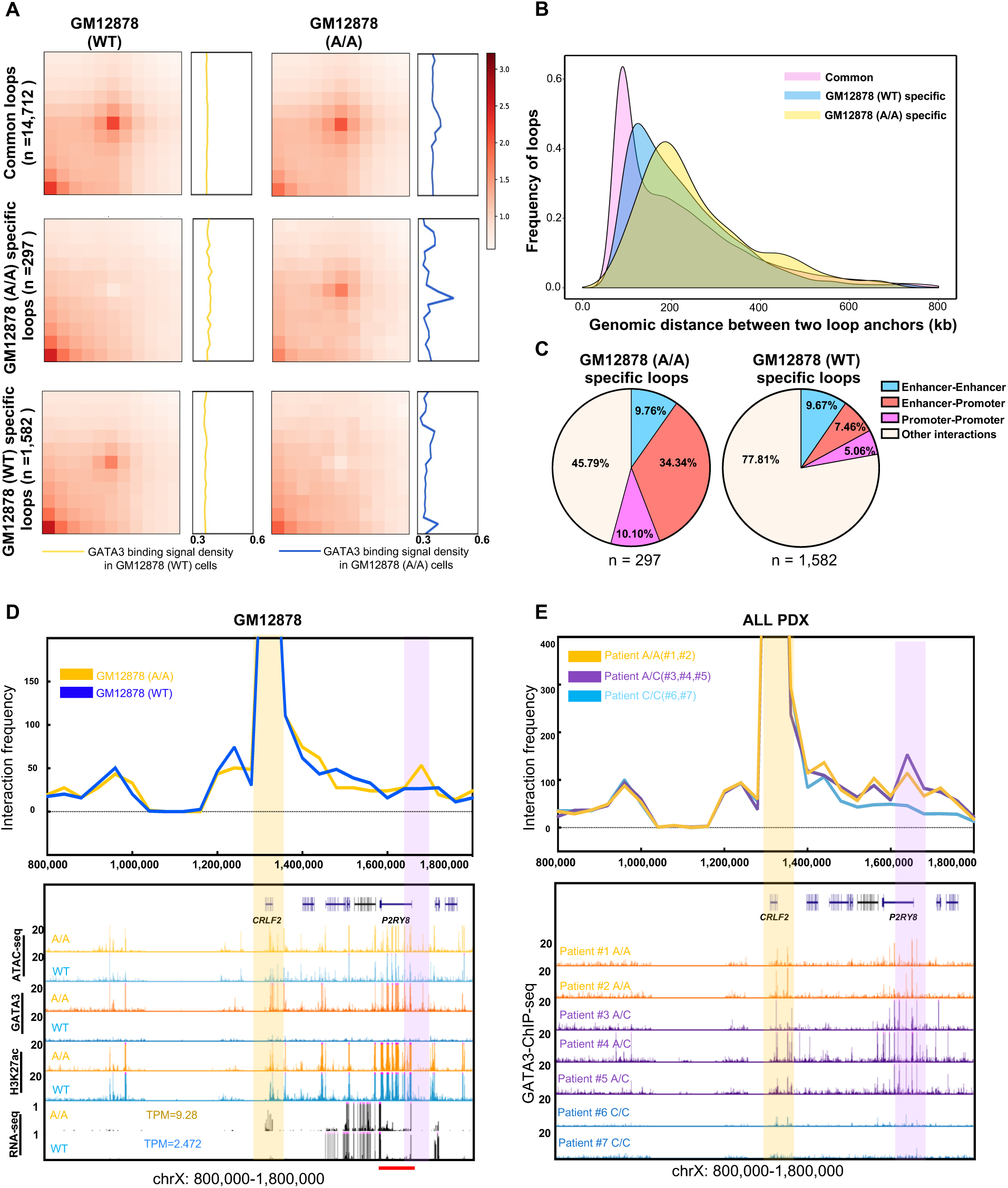
*GATA3* expression leads to new enhancer-promoter interactions, particularly in genes related to Ph-like ALL. **A**, APA plot indicates that GATA3 binding are enriched in engineered GM12878 (A/A) cell specific chromatin loops. **B**, Distance distribution of chromatin loops specific to GM12878 (A/A), GM12878 (WT), or common in both cell lines. **C**, Enhancer-Promoter and Promoter-Promoter are more enriched in the differential loops of engineered GM12878 (A/A) cells. **D-E**, Virtual 4-C analysis (40kb resolution) shows there is a A/A genotype-specific chromatin looping between the *P2RY8* enhancer (pink bar) and the *CRLF2* promoter (yellow bar) in engineered GM12878 (A/A) cells and also ALL PDX samples with A/A genotype. Red bar indicates the *P2RY8* super enhancer predicted by ROSE.

Inspired by this observation, we performed motif analysis of all the common breakpoint regions in Ph-like ALL patients (Roberts et al., 2014), and we observed an enrichment of GATA3 motif (**Figure S13A**). Finally, we examined the GATA3 ChIP-seq signals surrounding the Ph-like breakpoints in both the GATA3-overexpressed Nalm-6 ALL cells and engineered GM12878 cells, and again we observed an enrichment of GATA3 binding (**Figure S13B and S13C**). Taken together, these data provided evidence that GATA3 may be involved in chromosomal translocations in Ph-like ALL.

### *GATA3* directly regulates *CRLF2* pathways and contributes to the pathogenesis of Ph-like ALL

When ectopically expressed in ALL cell lines, *GATA3* induced a gene expression pattern that overlaps with the expression signature of Ph-like ALL(Perez-Andreu et al., 2013). In particular, inducible overexpression of *GATA3* led to up-regulation of *CRLF2* in a time-dependent manner (**Figure 5A**), with concomitant gain of GATA3 binding at the *CRLF2* promoter region overlapping with *CRLF2* rearrangement hotspots observed in Ph-like ALL (**Figure S14**). Conversely, down-regulation of *GATA3* by shRNA suppressed *CRLF2* transcription (**Figure 5B**), further indicating that GATA3 functions as a transcriptional regulator of *CRLF2*. It has been shown that CRLF2-mediated constitutive activation of the JAK-STAT pathway is responsible for leukemogenesis in hematopoietic cells(Mullighan et al., 2009). Therefore, we hypothesized that *GATA3* acts upstream of *CRLF2*, and the germline *GATA3* variant can directly influence CRLF2-JAK signaling (by upregulating *GATA3* expression). To test this possibility, we examined the effects of GATA3 on *in vitro* transforming potential and JAK-STAT signaling in mouse hematopoietic cell Ba/F3. *GATA3* overexpression resulted in upregulation of *CRLF2* and also led to phosphorylation of Jak2 and Stat5 (**Figure 5C**). Co-expression of *GATA3* and *JAK2^R683G^* were sufficient to induce cytokine-independent growth and Ba/F3 cell transformation, in a fashion analogous to co-expression of *CRLF2* and *JAK2^R683G^* although with a longer latency (**Figure 5D**). Interestingly, the addition of CRLF2 ligand, TSLP, potentiated transforming effects of GATA3 in Ba/F3 cells expressing mouse Il7r (Ba/F7 cells, **Figure S15**). These results strongly suggested that GATA3 directly up-regulates CRLF2 and thus impinges upon the pathogenesis of Ph-like ALL (**Figure 5E**).

**Figure 5.**
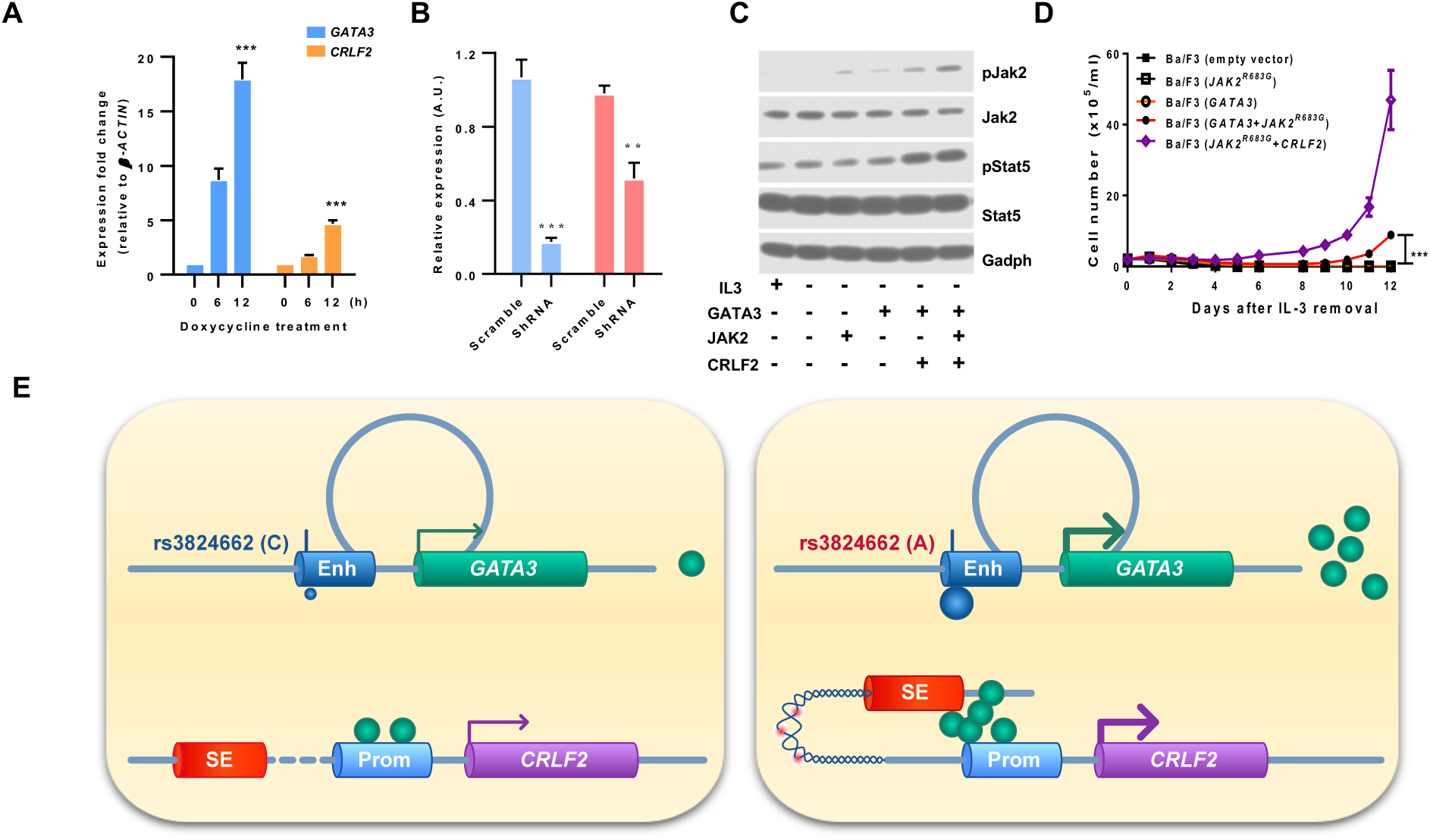
GATA3 potentiates CRLF2-JAK-STAT signaling in hematopoietic cells. **A-B**, *GATA3* regulates *CRLF2* transcription ALL cell line Nalm6 (overexpression in **A** and knockdown in **B**). The T bars indicate standard deviations (*p* value < 0.001 by 2-way ANOVA). **C**, JAK-STAT activation by GATA3. Mouse hematopoietic cell Ba/F3 was transduced with combinations of *GATA3*, *JAK2^R683G^*, and *CRLF2* as indicated, and cultured in the presence or absence of IL3. Phosphorylation of JAK2 and STAT5 was examined by immunoblotting with GAPDH as the loading control. **D**, IL3-independent growth of Ba/F3 cells transduced with *GATA3* alone, *JAK2^R683G^* alone, *GATA3* with *JAK2^R683G^*, *JAK2^R683G^* with *CRLF2*, or empty vector control. All the experiments were performed in triplicates (*p* value < 0.001 by 2way ANOVA). **E**, A schematic of our proposed model of how *GATA3* rs3824662 variant contributes to pathogenesis of Ph-like ALL. Risk “A” allele induces *GATA3* expression which binds to the *CRLF* promoter, loops *CRLF2* promoter to the super enhancer localized in *P2RY8* region, eventually resulting in *CRLF2* overexpression. The chromatin region between *CRLF2* promoter and *P2RY8* super enhancer also becomes more open and thus susceptible to damage (e.g., rearrangements). Enh: enhancer; SE: super enhancer; Prom: promoter.

## Discussion

Both inherited germline and somatic genetic variations contribute to the pathogenesis of different malignancies, including leukemias. Somatic genomic aberrations, i.e., mutations, rearrangements, insertion/deletion, have been shown to drive overt leukemogenesis by promoting the survival and proliferation of pre-leukemia hematopoietic cells. However, the roles of inherited leukemia risk variants, especially those in intronic/intergenic loci, remain largely unclear. For example, GWAS studies have identified 9 genomic loci with common SNPs associated with susceptibility to childhood ALL, but there has been little progress to move from descriptive association studies to identifying causative mechanisms relating these variants to ALL pathogenesis.

Here we define the regulatory function of a non-coding SNP rs3824462 associated with Ph-like ALL(Perez-Andreu et al., 2013). This variant strongly influences the susceptibility to high-risk ALL and also prognosis, consistently across different ALL treatment regimens(Migliorini et al., 2013; Perez-Andreu et al., 2013). In this work, we first reported that the rs3824662 variant is located inside an enhancer element and the risk allele showed significantly increased enhancer activity. Introducing the risk A allele at rs3824662 by CRSPR/Cas9 editing in the wildtype GM12878 cells directly confirmed its enhancer effects on *GATA3* transcription. Using a variety of chromatin conformation capturing techniques, we further demonstrated that this variant significantly reshaped chromatin interactions both locally and also in a global fashion. A recent study showed that GATA3 can act as a pioneer factor in the course of cellular reprogramming, making previously condensed chromatin more accessible by recruiting BRG1, a chromatin remodeling factor(Takaku et al., 2016). Similarly, our ATAC-seq data also suggested that the C-to-A allele substitution at rs3824662 resulted in many newly-gained open chromatin regions enriched for GATA3 binding sites, coupled with global 3D geome re-orginazation. In particular, we observed hundreds of regions switched from the active and open compartment to the repressive and compacted compartment. Among them are many essential genes whose expression are altered in Ph-like ALL, likely due to the change of chromatin environment. We also performed ATAC-seq, GATA3 ChIP-seq and Hi-C in a panel of seven ALL patient samples with different genotypes at rs3824462. In these analyses, we identified similar B-to-A switching in *PON2* genes and novel looping events betweeen *CRLF2* and *P2RY8* locus, indicating that these transcriptional regulation mechanisms are indeed operative in Ph-like ALL patients. However, these human leukemia samples harbor a plethora of somatic genomic abnoralities which likely confounded the effects from germline *GATA3* polymorphisms.

More interestingly, we found many GATA3 binding sites are located near the breakpoints of translocation events observed in Ph-like ALL, suggesting its over-expression might be related with chromosomal instability and susceptiblity to translocations. Therefore, we hypothesize that *GATA3* over-expression might facilitate enhancer hijacking, where a distal enhancer is rearranged to the proximity of oncogenes and leads to oncogenesis without gene fusions(Groschel et al., 2014; Hnisz et al., 2016; Northcott et al., 2014; Weischenfeldt et al., 2017). To further explore the role of GATA3 in genome instability, we also explored its binding profile in breast cancer, as GATA3 abnormal expression has also been reported in certain human breast cancer subtypes. We confirmed that GATA3 binding is also enriched in a breast cancer cell line (T47D) translocation breakpoints region as well (**Figure S16A**). Moreover, we also observed GATA3 and BRG1 co-localize at these translocation breakpoints (**Figure S16B and S16C**), suggesting potential intricate interactions between GATA3, BRG1 and genome instability.

Aberrantly high *GATA3* expression has been also identified in other B cell malignancies, such as classical Hodgkin lymphoma. Constitutive activation of NFkB and Notch-1 leads to higher *GATA3* expression in Reed Sternberg cells, which then contributes to cytokine secretion (especially IL13) and signaling typical in Hodgkin lymphoma(Stanelle et al., 2010). In contrast, *GATA3* is not expressed in normal B cells and in fact functions as a key regulator of lymphoid cell lineage commitment (B vs T cells)(Banerjee et al., 2013). The data we present in the current study points to novel roles of *GATA3* in global cellular reprogramming and pathogenesis of B-cell malignancies.

In conclusion, we report here that the inherited genetic variant rs3824662 is a cis-acting enhancer variant associated with *GATA3* transcription activation, which contributes to Ph-like ALL leukemogenesis through regulating *CRLF2* signaling. Our results suggest that transcription factor-mediated epigenomic reprograming can directly influence oncogene activity, and may be an important mechanism by which germline genetic variants influence cancer risk.

## Methods

### Patients

In this study, 5,008 childhood ALL patients were enrolled on Children’s Oncology Group (AALL0232(Larsen et al., 2016) and COG9904/9905/9906(Borowitz et al., 2008)) and St. Jude Children’s Research frontline clinical trials(Pui et al., 2010). Germline DNA was extracted from bone marrow samples or peripheral blood obtained from children with ALL during remission. This study was approved by institutional review boards at St. Jude Children’s Research Hospital and COG affiliated institutions and informed consent was obtained from parents, guardians, or patients, as appropriate. Ph-like ALL status was determined on the basis of global gene expression, as described previously(Perez-Andreu et al., 2013). Patient-derived xenograpts of ALL were selected from the St. Jude PROPEL resource with genomic characterization and sample authentication described at https://stjuderesearch.org/site/data/propel

### *GATA3* targeted sequencing

Illumina dual-indexed libraries were created from the germline DNA of 5,008 children with ALL and pooled in sets of 96 before hybridization with customized Roche NimbleGene SeqCap EZ probes (Roche, Roche NimbleGen, Madison, WI, USA) to capture the *GATA3* genomic region. Quantitative PCR was used to define the appropriate capture product titer necessary to efficiently populate an Illumina HiSeq 2000 flow cell for paired-end 2 × 100 bp sequencing. Coverage of greater than 20 x depth was achieved across more than 80% of the targeted regions for nearly all samples. Sequence reads in the FASTQ format were mapped and aligned using the Burrows-Wheeler Aligner (BWA)(Li and Durbin, 2009), and genetic variants were called using the GATK pipeline (version 3.1)(Poplin et al., 2017), as previously described, and annotated using the ANNOVAR(Wang et al., 2010) program with the annotation databases including RefSeq(O’Leary et al., 2016). All the *GATA3* non-silent variants were manually reviewed in the Integrative Genomics Viewer(Robinson et al., 2011). Association of genotype with Ph-like ALL status was examined following our stablished statistical procedure(Perez-Andreu et al., 2013), i.e., comparing allele frequency in ALL cases with vs without the Ph-like gene expression signature, using the logistic regression test with genetic ancestry as covariables.

### Knock-in rs3824662 risk allele in GM12878

sgRNA targeted rs3824662 locus was cloned into CRISPR-CAS9 vector PX458 (Addgene)(Ran et al., 2013) and co-transfected into GM12878 along with single-strand donor DNA which carries risk allele A (**Supplementary Table 2**). After 68h of transfection, GFP positive cells were sorted into 20 96-well plates (color BD FACS Aria SORP high-performance cell sorter). Half of cells from successfully expanded clones were transferred into 24-well plates and the genomic DNA of the rest cells was extracted for PCR rs3824662 region. Pst1 (NEB) restriction enzyme digestion was used to select the heterozygous or homozygous knock-in clones. Successful knock-in clones were confirmed by Sanger sequencing.

### 3D chromatin structure mapping by Hi-C

Hi-C in GM12878 cells and PDX samples were performed using the Arima-HiC kit as per the manufacturer’s instructions. Briefly, 1 million GM12878 WT, A/A cells and PDX sample were fixed with 1% formaldehyde, digested with restriction enzyme, end-labeled with Biotin-14-dATP, and then followed by ligation. The ligated chromatin was reverse-crosslinked and sonicated by Covaris E220 to produce 300–500 bp fragments. Biotin labeled DNA fragments were isolated using dynabeads Streptavidin C1 beads and followed by end-repair, adenylation, adaptor ligation and PCR amplification. The quantity of the library was measured by both BioAnalyzer (Agilent) and Kapa Library Quantification Kit (Kapa Biosystems). Finally, the library was performed pair-end 2×100bp high-throughput sequencing using HiSeq 2500 and Nova-seq (Illumina).

### Cytokine-dependent growth assay in Ba/F3 cells and Ba/F7 cells

The full-length *GATA3* and *CRLF2* coding sequence were purchased from GE Healthcare and cloned into the cL20c-IRES-GFP lentiviral vector. cl20c-*CRLF2*-IRES-GFP was modified into cl20c-*CRLF2*-IRES-CFP, and lentiviral supernatants were produced by transient transfection of HEK-293T cells using calcium phosphate. The MSCV-JAK2^R683G^-IRES-GFP construct was a gift from Dr. Charles Mullighan at St. Jude Children’s Research Hospital(Mullighan et al., 2009) and modified into MSCV-JAK2R683G-IRES-mCherry and retroviral particles were produced using 293T cells. Ba/F3 cells and Ba/F7 cells were maintained in medium supplemented with 10 ng/ml recombinant mouse interleukin 3 (IL3) and interleukin 7 (IL7) (PeproTech), respectively. Ba/F3 or Ba/F7 cells were transduced with lentiviral supernatants expressing *GATA3*. GFP positive cells were sorted 48 hours after *GATA3* transduction and maintained in the IL3 medium for another 24 hours before transfected by *JAK2^R683G^* retroviral supernatants. Forty-eight hours later, GFP/mCherry double positive cells were sorted and maintained in medium with respective cytokine for 48 hours. Cells transduced with empty vector, *JAK2 ^R683G^* or *JAK2^R683G^* and *CRLF2* were sorted out for controls. Then, cells were washed three times and grown in the absence of cytokine. For TSLP assay, cells are maintained in medium with 10 ng/ml TSLP but without IL3. Cell growth and viability were monitored daily by Trypan blue using a TC10 automated cell counter (BIO-RAD). Each experiment was performed three times.

Additional experimental details and data analyses are included in the **Supplementary Methods**.

## Acknowledgements

This work was supported by the US National Institutes of Health (CA21765, CA98543, CA114766, CA98413, CA180886, CA180899, GM92666, GM115279, and GM097119) and the American Lebanese Syrian Associated Charities. H.Z. is a St. Baldrick’s International Scholar (grant 522589) and supported by the National Science Foundation of China (81300401). S.P.H. is the Jeffrey E. Perelman Distinguished Chair in Pediatrics at The Children’s Hospital of Philadelphia. M.L.L. is the UCSF Benioff Chair of Children’s Health and the Deborah and Arthur Ablin Chair of Pediatric Molecular Oncology. F.Y. is supported by 1R35GM124820, R01HG009906, U01CA200060 and R24DK106766. We thank the patients and parents who participated in the St. Jude and COG clinical trials included in this study, the clinicians and research staff at St Jude Children’s Research Hospital and COG institutions.

## Author Contributions

The study was conceived by JJ.Y. and F.Y., designed by JJ.Y., F.Y., H.Y. and H.Z., and supervised by JJ.Y. and F.Y.. H.Y. and T.L. performed the CRISPR Knock-in, Hi-C, ChIP-seq and ATAC-seq experiments in GM12878 and patient PDX samples under F.Y.’s supervision. H.Z. performed targeted-resqeuencing in cohorts and leukemia transforming assay in Ba/F3 and Ba/F7 cells. Data preprocessing was conducted by Y.L., M.Q., B.Z.,W.Y and H.Y.; statistical analyses by Y.L., H.Y. and H.Z.; data interpretation by JJ.Y., F.Y., H.Y., H.Z., Y.L., T.L., M.Q., B.Z., Y.L., J.X., I.S., W.Y., KG. R., V.P-A., H.X., J. G-F., C.S., C-H.P., W.E.E., M.V.R., S.P.H., C.G.M. and M.L.L. JJ.Y., F.Y., H.Y. and H.Z. wrote the manuscript; All authors approved the final version for publication.

## Supplementary Methods

### Functional studies

#### Luciferase reporter gene assay

A 1,120-bp region encompassing the *GATA3* SNP rs3824662 was amplified using CloneAmp HiFi PCR Premix (Clontech) (primer sequences in **Supplementary Table 4**) and then cloned into the pGL4.23-mini/P vector with a minimal SV40 promoter upstream of the firefly luciferase gene sequence. For reporter assays, 2×10^6^ SUP-B15, GM12878, Ba/F3 cells were resuspended in 100 μl of Nucleofector Solution Kit V (Lonza) with the addition of 1.9 μg of pGL4.23 constructs and 100ng of renilla pTK plasmid. Cells were electroporated and then incubated at 37°C for 24 hours with 5% CO_2_. Similarly, HEK293T cells (6×10^4^) were plated on 96-well plate (flat bottom), and co-transduced with 95 ng pGL4.23 constructs and 5ng renilla pTK, and then incubated for 24 hrs. Luciferase activity was measured using the Dual-Glo Luciferase Assay system (Promega). Experiments were performed in triplicate. To control for cell number and transfection efficiency, firefly luciferase activity was normalized to renilla luciferase. Measurements were presented as a ratio relative to the activity of the pGL4.23-mini/P empty vector.

#### Inducible *GATA3* overexpression and *GATA3* knockdown

Full-length *GATA3* cDNA was cloned into the lentiviral vector pLV-tetON (a gift from Dr. Chunliang Li at St. Jude Children’s Research Hospital). Nalm6 and SUP-B15 cells were transduced with pLV-*GATA3*-tetON for 48 hours and then subjected to bleomycin selection (0.5 mg/ml). Single clones were established in which doxycycline-induced *GATA3* overexpression was confirmed by RT-qPCR and immuno-blotting.

The lentiviral pLKO.1 constructs with *GATA3* shRNA and scrambled shRNA were purchased from Sigma-Aldrich. Nalm6 and SUP-B15 cells were lentivirally transduced with pLKO.1-*GATA3*-shRNA or scrambled shRNA for 48 hours and then subjected to puromycin selection (1.0 ug/ml). The degree of *GATA3* and *CRLF2* knockdown was evaluated by RT-qPCR.

#### Jak-Stat activation

In transduced Ba/F3 cells, Jak-Stat pathway activation was evaluated by immunoblotting using anti-Jak2 antibody (Cell Signaling, 3230, 1:1000 dilution), anti-phospho-Jak2 antibody (Cell Signaling, 3771, 1:1,000 dilution), anti-Stat5 antibody (Cell Signaling, 9310, 1:1,000 dilution), and anti-phosphor-Stat5 antibody (Cell Signaling, 9314, 1:1,000 dilution). Gadph was used as a loading control.

### Transcriptomic and epigenomic profiling

#### RNA-seq

Total RNAs were extracted from 5 million engineered GM12878 cells using Trizol (Invitrogen). cDNA libraries were prepared using SureSelect Strand Specific RNA Library Preparation Kit (Agilent). Briefly, polyA RNA was purified from 1000 ng of total RNA using oligo(dT) beads (Invitrogen) and then fragmented, followed by reverse transcription, end repair, adenylation, adaptor ligation and subsequent PCR amplification. The final product was checked by size distribution and concentration using BioAnalyzer High Sensitivity DNA Kit (Agilent) and Kapa Library Quantification Kit (Kapa Biosystems). Pair-end 2×50bp high-throughput sequencing was performed using HiSeq 2500 (Illumina). Expression of *GATA3*, *CRLF2*, *KIN*, *ENSG*, *SFMBT2*, *ITIH5*, *ITIH2*, *FAF3*, and *GATA*-*AS* was also quantified by qRT-PCR with ACTIN as loading control (Primer sequences of indicated genes were listed in **Supplementary Table 4**).

#### ATAC-seq

ATAC-seq was performed as previously described(Buenrostro et al., 2015). A total of 30,000-50,000 live cells were collected, washed once in PBS and resuspended in 50 ul ATAC-seq lysis buffer (10 mM Tris-HCl pH 7.4, 10 mM NaCl, 3 mM MgCl_2_ and 0.1% IGEPAL CA-630) followed by immediate centrifugation for 10 min at 4 °C. Cell pellets were then resuspended in 50 ul reaction solution which contained 1XTD buffer and 2.5 ul Tn5 transpose (Illumina, FC-121-1030) and incubated at 37 °C for 60 mins. The fragmented DNA was purified by MinElute kit (Qiagen) and amplified by PCR. The final product was checked by size distribution and concentration using BioAnalyzer High Sensitivity DNA Kit (Agilent). Pair-end 2×50bp high-throughput sequencing was performed using HiSeq 2500 (Illumina).

#### ChIP-seq and ChIP-qPCR

ChIP-seq and ChIP-qPCR were performed as previously described(Shen et al., 2012). Briefly, 100 ug chromatin was sonicated to 100-300 bp by Covaris E220, and 5 ug chromatin was used as input. Bead-antibody complex was prepared by incubating 11 ul of sheep anti-mouse IgG dynabeads (ThermoFisher, 11201D) or sheep anti-rabbit IgG dynabeads (ThermoFisher, 11203D) with 3ug of anti-H3K4me1 (Abcam, ab8895), anti-H3K27ac (Active motif, 39133), NF-1 (Santa Cruz, sc-74444), anti-GATA3 (Santa Cruz, sc268) or mouse IgG (ThermoFisher, 10400C), at 4°C for 4 hours with shaking. Then fragmented chromatin was incubated with bead-antibody complex overnight with shaking followed by stringent wash, elution and reverse crosslinking. For ChIP-seq, the immunoprecipitated DNA and input DNA were processed by end repair, adenylation, adaptor ligation, PCR amplification and subsequent size selection using AMPure XP beads (Beckman). 2×50 or 2×100 bp high-throughput sequencing procedures were performed using HiSeq 2500 (Illumina). For ChIP-qPCR, the measurements of target loci binding enrichment by specific antibody and mouse IgG were normalized to input DNA, respectively.

#### Capture-C

Capture-C was performed as previously described(Huang et al., 2017). Briefly, 10 million engineered or wildtype GM12878 cells were fixed with 1% formaldehyde and digested with DpnII (NEB), followed by DNA ligation. The ligated chromatin was then reverse-crosslinked, purified by phenol-chloroform and followed by sonication to produce 200– 300 bp fragments using Covaris E220. Fragmented DNA was used to make libraries with the NEBNext DNA Library Prep Master Mix Set (NEB). Hybridization with 60 bp biotinylated capture probes (**Supplementary Table 5**) was performed with the xGen® Lockdown® Reagents (Integrated DNA Technologies). In brief, 3C libraries were dried and resuspended with hybridization reagents. 3 pmol pooled capture probes was mixed with the resuspended libraries and incubated for 72 hr at 47°C. After streptavidin beads purification and PCR, the pulldown material was treated with a second round of 24-hr incubation to improve specificity. The capture probes, ordered from Integrated DNA Technologies, flank DpnII sites proximal to rs3824662.

### Data analyses

#### Sequencing QC

TrimGalore (https://github.com/FelixKrueger/TrimGalore) was used to trim and filter all the illumina next generation sequencing fastq reads, including ChIP-seq, RNA-seq, Capture-C and HiC with the following parameters: -q 20 --phred33 --paired --trim-n (**Supplementary Table 6-8**).

#### Capture C process

Capture C data were processed by Hi-C-Pro(Servant et al., 2015), we required one of the pairs should be mapped to the rs3824662 anchor regions. CHiCAGO(Cairns et al., 2016) was used to assign significant interactions linked to the captured rs3824662 fragments.

#### RNA-seq data process

STARv2.6.0(Dobin et al., 2013) was used to align RNA-seq data to female hg19 reference genomes (https://www.encodeproject.org/files/female.hg19/) with “outSAMtype BAM SortedByCoordinate-quantMode TranscriptomeSAM” parameters. The genome-wide signal coverage tracks were generated by using STARv2.5.3 with the “-outWigStrand Stranded” parameter. RSEM(Li and Dewey, 2011) (https://github.com/deweylab/RSEM) was used to quantify and calculate the expression values for known genes (https:// www. gencodegenes.org/human/release_19.html). Genes with TPM value less than 1 in all samples were removed. DEseq2(Love et al., 2014) were used to identify differential expression genes with *p*-value<0.05 and log_2_Fold-change>2 as the cut-off. The GO term of differential expression genes was analyzed by Panther(http://pantherdb.org/).

#### ChIP-seq data process

Pair-end sequencing data were mapped to the female reference genome (hg19) using bowtie2(version 2.3.4.3)(Langmead and Salzberg, 2012). Non-uniquely mapping reads (MAPQ<30) were removed, and PCR duplicate reads were removed by Picard (http://broadinstitute.github.io/picard/). ENCODE-chip-seq-pipeline2 were followed to call the peaks, the narrow peaks with Poisson *p*-value greater than 0.001 were removed to ensure good quality peaks for further analysis. To further qualify the predicted peaks, Reads Per Million (RPM) of IP data and input data in each peak region were calculated and the qualified peaks should pass the threshold of two-fold enrichment (RPM_IP_/RPM_input_>2) and RPM_IP_-RPM_input_>1. To check repeatability between biological replicates, firstly we divided the reference genomes into 10kb bins and computed the number of reads within each bin. The Pearson correlation coefficient between each biological replication was calculated using above-normalized 10kb bins reads. IDR with a threshold of 0.05 was used to measure the reproducibility of peaks from replicates(Li et al., 2011). The peaks were re-centered and set to a fixed width of 250 bp and identified differential GATA3 peaks using the DiffBind R package(Ross-Innes et al., 2012), The genome-wide ChIP-seq signal tracks were generated by MACS2(V2.2.4)(Zhang et al., 2008) for TFs and histone marks.

#### The GWAS hints enrichment analysis

The GWAS hints were download from https://www.ebi.ac.uk/gwas/. 10,279 disease-associated and at least identified by 2 articles single-nucleotide polymorphisms were selected. Investigated their distribution at differential GATA3 peaks 10kb flanking regions.

#### ATAC-seq data process

Pair-end sequencing data were mapped to the female reference genome (hg19) using bowtie2(version 2.3.4.3)(Langmead and Salzberg, 2012) with -X 2000 parameter. Non-uniquely mapping reads (MAPQ<30) were removed, and PCR duplicate reads were removed by Picard (http://broadinstitute.github.io/picard/). ENCODE-atac-seq-pipeline were followed to call the peaks, the narrow peaks with Poisson p-value greater than 0.001 were removed to ensure good quality peaks for further analysis. To check repeatability between biological replicates, firstly we divided the reference genomes into 10kb bins and computed the number of reads within each bin. The Pearson correlation coefficient between each biological replication were calculated using above-normalized 10kb bins reads. IDR with a threshold of 0.05 was used to measure the reproducibility of peaks from replicates(Li et al., 2011). The genome-wide ATAC-seq signal tracks were generated by MACS2(V2.2.4)(Zhang et al., 2008) for TFs and histone marks. ATAC-seq peaks (called by MACS2 using parameters –nomodel --broad --keep-dup all -shift-75 --extsize 150) was merged with a 110 bp-window. Nucleosome position with in these peak regions were then called using the NucleoATAC software(Schep et al., 2015) (https://github.com/GreenleafLab/NucleoATAC) version 0.3.4 with default parameters, and normalized nucleosome occupancy signal value was used to plot the nucleosome position profile. Footprint was identified by the HINT software (Hmm-based IdeNtification of Transcription factor footprints)(Li et al., 2019) based on ATAC-seq data. Briefly, ATAC-seq narrowpeaks were used as input, the footprint region were filtered by footprint score>10, transcription factor motifs overlap with footprints was identified using the MOODS package (https://github.com/jhkorhonen/MOODS)(Korhonen et al., 2009), with motifs from the HOCOMOCO database (http://hocomoco11.autosome.ru/)(Kulakovskiy et al., 2018).

#### Hi-C data process

The Hi-C data were aligned to the female reference genome(hg19) by bwa mem model(Li and Durbin, 2009) with -SP5M parameters. The PCR duplications and low-quality aligned pairs were removed by pairtools (https: //github.com/ mirnylab/pairtools), the “UU”, “UR” and “RU” types pair were kept for further analysis. We generated 5kb, 10kb, 25kb, 40kb,50kb,100kb muti-resolutions balanced cool file and hic file for visualization. Correlations between Hi-C replicates were calculated HiCRep(Yang et al., 2017). We combined biological replicates of Hi-c data from each engineered and GM12878 clone.

**A and B compartments** were identified using previously described(Lieberman-Aiden et al., 2009) with modifications. We construct raw 10kb Hi-C contact matrix without normalization of each cell type and patient, then calculated the expected interaction frequency between any two bins given the distance separating them in the genome. The observed/expected matrix was generated and then converted to a Pearson correlation matrix. Principal component analysis is applied to the correlation matrix similar as previously described. The value on first principal component for each bin was used to correlate with ATAC-seq signal to assign a genomic region to A or B compartment. If the sign of PC1 value changed between engineered GM12878 cell lines with different genotypes at rs3824662, we considered it as the A/B switch region.

**The insulation score** was calculated by the Perl script matrix2insulation.pl (Record Owner) at 40kb resolution matrix with “–ss 80000 --im iqrMean --is 480000 --ids 320000” parameters. The topologically associated domains were identified by the Perl script insulation2tads.pl, the 0.3 of min boundary strength was set as threshold.

**The interaction loops** were identified by Peakachu(Salameh et al., 2019) in 10kb resolutions, the models for predicting loops using H3K27ac 10% and CTCF 10% model. The predicted loops pass the probability score threshold great than 0.8. For the differential loops, we first calculated the probability score of each pair cross the genome, then we compare the probability score of predicted loops in sample A and probability score of pairs in sample B, we set 2-fold change as the cut off.

**The Virtual 4C plot** were used the embed method from the 3D genome browser(Wang et al., 2018). Briefly, a bait (for example, CRLF2 locus) and flanking region were chosen, then the row overlapping the bait and flanking regions were extracted from the Hi-C matrix. The number of observed contacts was plotted with a smoothing window to obtain virtual 4C profiles. To ensure the interaction frequency from different library comparable, the interactions in chromosome X were normalized by the number of interactions in viewpoints.

#### GATA3 and BRG1 binding in cancer breakpoints

Ph-like patient breakpoints and T47D cancer cell line breakpoints were collected from previous study(Dixon et al., 2017; Reshmi et al., 2017) and expanded to +/- 1kb region. Motif enrichment analysis was performed using HOMER version 4.8 findMotifsGenome.pl function for exploring potential transcription factor binding within these expanded breakpoint regions. GATA3 binding signal in these expanded breakpoint regions was generated by deeptools using GATA3 ChIP-seq in the following cells: GM12878 C/C clone, GM12878 A/A clone, Nalm6, Nalm6_gata3overexpressing and T47D breast cancer cell(Adomas et al., 2014). BRG1 ChIP-seq in T47D breast cancer cell were collected from GSE112491 are plotted the same way.

---

**Supplementary Table 1.** Clinical charateristics of ALL patients included in *GATA3* sequencing.

| Features | Group | Ph-like ALL | Non Ph-like ALL | Other B-ALL |
| --- | --- | --- | --- | --- |
|  |  | N=143 | N=852 | N=4013 |
| Age at diagnosis (%) | < 10 yrs | 48 (33.6) | 410 (48.1) | 2141 (53.4) |
|  | >= 10 yrs | 93 (65.0) | 434 (50.9) | 1351 (33.7) |
|  | NA | 2 ( 1.4) | 8 ( 0.9) | 521 (13.0) |
| Gender (%) | Female | 55 (38.5) | 393 (46.1) | 1837 (45.8) |
|  | Male | 88 (61.5) | 459 (53.9) | 2165 (53.9) |
|  | NA | 0 ( 0.0) | 0 ( 0.0) | 11 ( 0.3) |
| Leukocyte count at diagnosis (%) | < 50x109/L | 84 (58.7) | 525 (61.6) | 2497 (62.2) |
|  | >= 50x109/L | 57 (39.9) | 319 (37.4) | 996 (24.8) |
|  | NA | 2 ( 1.4) | 8 ( 0.9) | 520 (13.0) |
| Leukemia cell DNA index (%) | < 1.16 | 128 (89.5) | 684 (80.3) | 2606 (64.9) |
|  | >= 1.16 | 13 ( 9.1) | 161 (18.9) | 802 (20.0) |
|  | NA | 2 ( 1.4) | 7 ( 0.8) | 605 (15.1) |

**Supplementary Table 2.** Information of ALL PDXs used in this study.

| Patient ID | Xeno ID | Sample ID | Tumor Type | Tumor Subtype | Genotype | GATA3 Genotype | GATA3 Expression | Fusion | PDX mouse ID |
| --- | --- | --- | --- | --- | --- | --- | --- | --- | --- |
| #1 | PANZPJ | SJBALL020579 | B-ALL | Ph-like | 0 | aa | 11.24 | IGH-EPOR | 28, 29, 30 |
| #2 | PANWJB | SJBALL020589 | B-ALL | Ph-like | 0 | aa | 11 | ATF7IP-JAK2 | 22, 23, 24, 25, 26, 27 |
| #3 | TB-00-1196 | SJBALL021102 | B-ALL | Ph-like | 1 | ac | 7.99 | NoFusion | 40, 41, 42 |
| #4 | PARJCY | SJBALL020625 | B-ALL | Ph-like | 1 | ac | 9 | ZC3HAV1-ABL2 | 19, 20, 21 |
| #5 | PASMNW | SJBALL020980 | B-ALL | Ph-like | 1 | ac | 9.2 | PAG1-ABL2 | 16, 17, 18 |
| #6 | TB-07-1094 | SJPHAL L008 | B-ALL | BCR-ABL1 | 2 | cc | 5 | BCR-ABL1 | 31, 32, 33 |
| #7 | TB-04-2227 | SJBALL205 | B-ALL | NUTM1 | 2 | cc | 4.39 | CUX1-NUTM1 | 34, 35, 36 |

**Supplementary Table 3.**
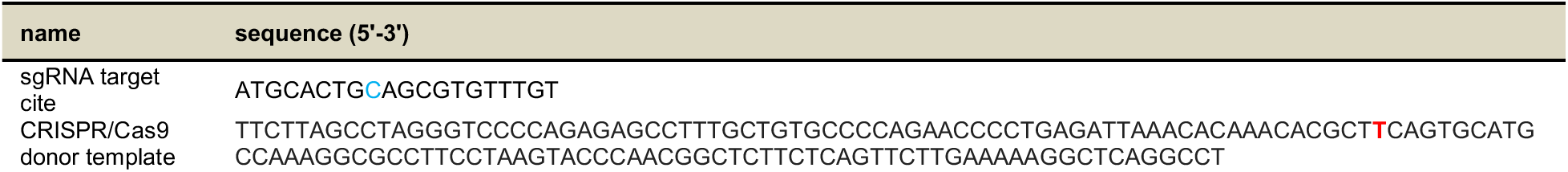
CIRPSR target sites and donor sequence for rs824662 knock-in in the GM12878 cell line.

**Supplementary Table 4.**
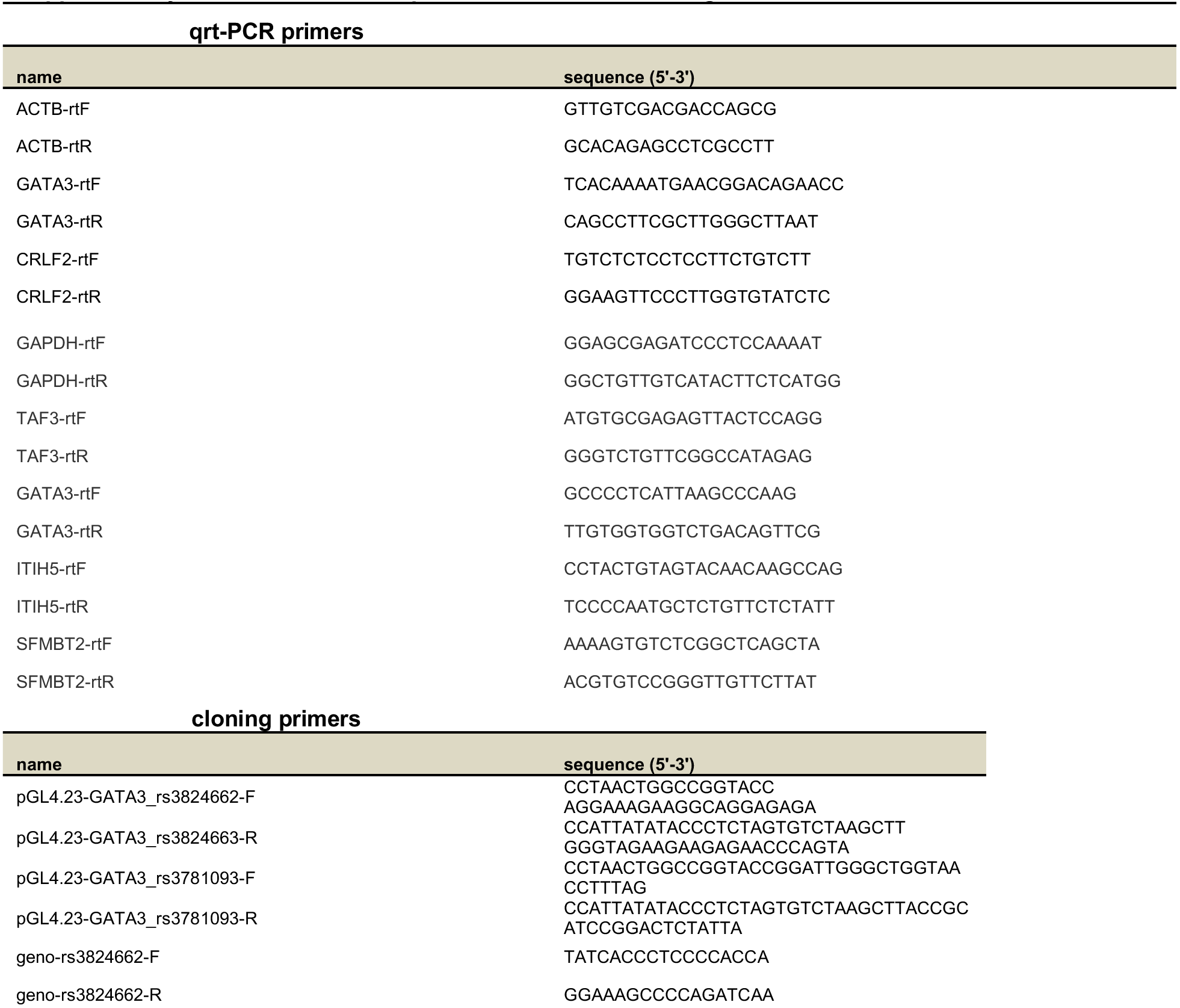
Primers for quantitative PCR and cloning.

**Supplementary Table 5.**
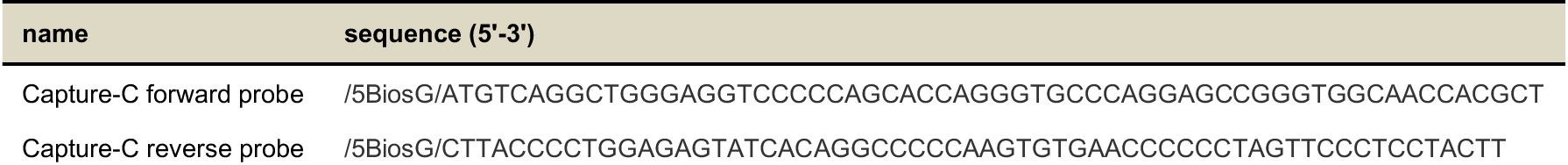
Probe sequence for capture C experiments.

**Supplementary Table 6.**
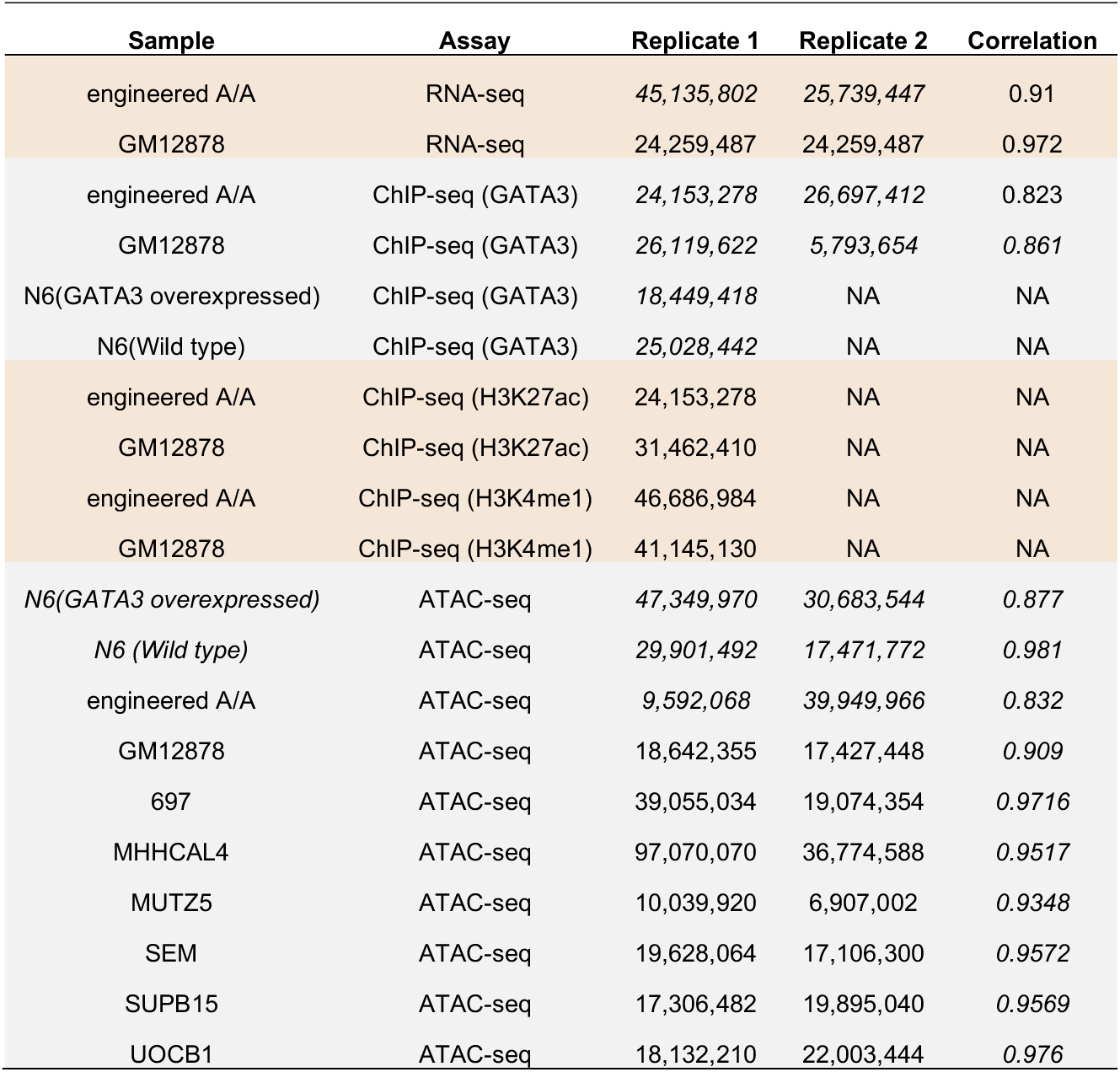
Reads number of RNA-seq, ATAC-seq and ChIP-seq in engineered GM12872 cell lines and human ALL cell lines.

| Sample | Assay | Replicate 1 | Replicate 2 | Correlation |
| --- | --- | --- | --- | --- |
| engineered A/A | RNA-seq | 45,135,802 | 25,739,447 | 0.91 |
| GM12878 | RNA-seq | 24,259,487 | 24,259,487 | 0.972 |
| engineered A/A | ChIP-seq (GATA3) | 24,153,278 | 26,697,412 | 0.823 |
| GM12878 | ChIP-seq (GATA3) | 26,119,622 | 5,793,654 | 0.861 |
| N6(GATA3 overexpressed) | ChIP-seq (GATA3) | 18,449,418 | NA | NA |
| N6(Wild type) | ChIP-seq (GATA3) | 25,028,442 | NA | NA |
| engineered A/A | ChIP-seq (H3K27ac) | 24,153,278 | NA | NA |
| GM12878 | ChIP-seq (H3K27ac) | 31,462,410 | NA | NA |
| engineered A/A | ChIP-seq (H3K4me1) | 46,686,984 | NA | NA |
| GM12878 | ChIP-seq (H3K4me1) | 41,145,130 | NA | NA |
| N6(GATA3 overexpressed) | ATAC-seq | 47,349,970 | 30,683,544 | 0.877 |
| N6 (Wild type) | ATAC-seq | 29,901,492 | 17,471,772 | 0.981 |
| engineered A/A | ATAC-seq | 9,592,068 | 39,949,966 | 0.832 |
| GM12878 | ATAC-seq | 18,642,355 | 17,427,448 | 0.909 |
| 697 | ATAC-seq | 39,055,034 | 19,074,354 | 0.9716 |
| MHHCAL4 | ATAC-seq | 97,070,070 | 36,774,588 | 0.9517 |
| MUTZ5 | ATAC-seq | 10,039,920 | 6,907,002 | 0.9348 |
| SEM | ATAC-seq | 19,628,064 | 17,106,300 | 0.9572 |
| SUPB15 | ATAC-seq | 17,306,482 | 19,895,040 | 0.9569 |
| UOCB1 | ATAC-seq | 18,132,210 | 22,003,444 | 0.976 |

**Supplementary Table 7.** Reads number of RNA-seq, ATAC-seq and ChIP-seq in ALL PDX samples.

| Sample | Assay | Replicate 1 |
| --- | --- | --- |
| PDX17_A-C_SJBALL020980 | RNA-seq | 146,360,710 |
| PDX19_A-C_SJBALL020625 | RNA-seq | 103,250,506 |
| PDX23_AA_SJBALL020589 | RNA-seq | 125,583,968 |
| PDX29_AA_SJBALL020579 | RNA-seq | 197,024,688 |
| PDX31_CC_SJPHALL008 | RNA-seq | 61,401,228 |
| PDX34_CC_SJBALL205 | RNA-seq | 127,885,146 |
| PDX40_AC_SJBALL021102 | RNA-seq | 101,083,624 |
| PDX17_A-C_SJBALL020980 | ATAC-seq | 26,470,800 |
| PDX19_A-C_SJBALL020625 | ATAC-seq | 29,299,466 |
| PDX23_AA_SJBALL020589 | ATAC-seq | 30,213,231 |
| PDX29_AA_SJBALL020579 | ATAC-seq | 27,725,249 |
| PDX31_CC_SJPHALL008 | ATAC-seq | 27,725,249 |
| PDX34_CC_SJBALL205 | ATAC-seq | 30,043,076 |
| PDX40_AC_SJBALL021102 | ATAC-seq | 30,202,445 |
| PDX17_A-C_SJBALL020980 | ChIP-seq | 26,524,384 |
| PDX19_A-C_SJBALL020625 | ChIP-seq | 27,539,028 |
| PDX23_AA_SJBALL020589 | ChIP-seq | 15,944,315 |
| PDX29_AA_SJBALL020579 | ChIP-seq | 13,415,639 |
| PDX31_CC_SJPHALL008 | ChIP-seq | 14,790,057 |
| PDX34_CC_SJBALL205 | ChIP-seq | 14,805,006 |
| PDX40_AC_SJBALL021102 | ChIP-seq | 13,378,167 |

**Supplementary Table 8.** Reads number of Hi-C libraries in GM12878 and ALL PDX samples.

| Sample | Assay |  | useful_reads | cis_interaction | tran_interaction | cis_interaction>20kb | correlation |
| --- | --- | --- | --- | --- | --- | --- | --- |
| engineered A/A (49) | Hi-C | Rep1 | 309,055,073 | 221,422,151 | 87,632,922 | 172,346,444 | 0.926 |
| engineered A/A (7) | Hi-C | Rep2 | 238,327,534 | 238,327,534 | 67,632,938 | 133,937,841 |  |
| GM12878 | Hi-C | Rep1 | 461,848,818 | 359,704,885 | 102,143,933 | 257,627,006 | 0.911 |
| GM12878 | Hi-C | Rep2 | 104,779,887 | 81,752,318 | 23,027,509 | 58,039,429 |  |
| PDX17_A-C_SJBALL020980 | Hi-C | Rep1 | 278,196,257 | 229,769,937 | 48,426,320 | 159,128,729 |  |
| PDX19_A-C_SJBALL020625 | Hi-C | Rep1 | 249,173,169 | 216,029,066 | 33,144,103 | 150,159,056 |  |
| PDX23_AA_SJBALL020589 | Hi-C | Rep1 | 621,592,890 | 545,596,312 | 75,996,578 | 332,074,940 |  |
| PDX29_AA_SJBALL020579 | Hi-C | Rep1 | 475,527,653 | 412,460,038 | 63,067,615 | 251,809,325 |  |
| PDX31_CC_SJPHALL008 | Hi-C | Rep1 | 399,989,579 | 350,354,228 | 49,635,351 | 238,734,093 |  |
| PDX34_CC_SJBALL205 | Hi-C | Rep1 | 582,808,716 | 508,366,206 | 74,442,510 | 343,373,212 |  |
| PDX40_AC_SJBALL021102 | Hi-C | Rep1 | 393,681,539 | 335,905,716 | 57,775,823 | 228,952,192 |  |

**Supplementary Figure 1.**
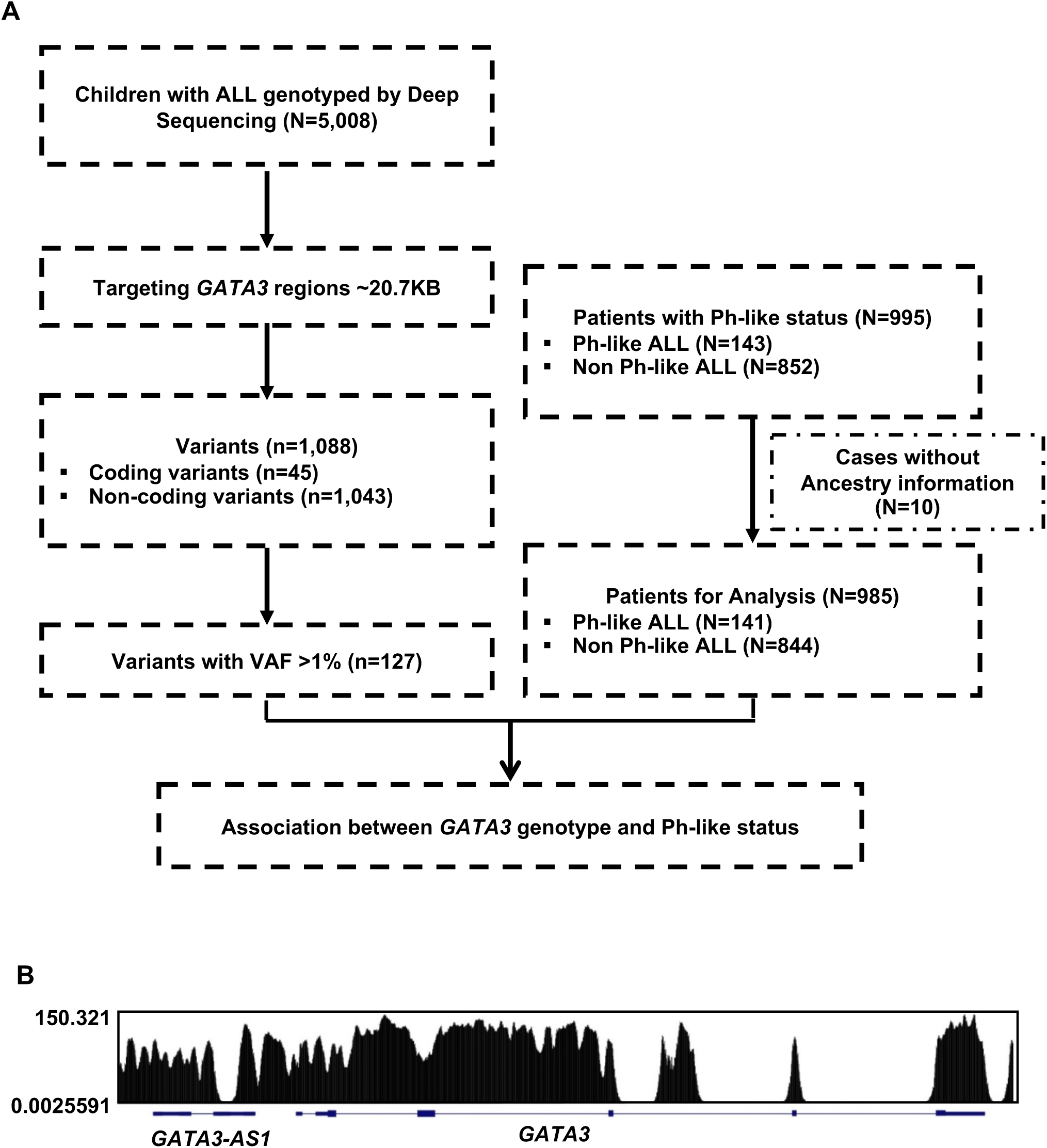
Targeted *GATA3* sequencing in 5,008 children with ALL. **A**, Flow chart of Ph-like ALL risk variant discovery. *GATA3* variants were identified from 5,008 children with ALL, of whom 995 patients were examined for Ph-like subtype (143 Ph-like *vs*. 852 non-Ph-like ALL). A total of 127 variants with sufficient frequency were subjected to association test in this subset. **B**, Read density and coverage of the *GATA3* target region. We covered coding region, 3kb upstream of 5’UTR, 1kb after 3’UTR, and all predicted open chromatin regions (based on ATAC-seq data in GM12878 cells).

**Supplementary Figure 2.**
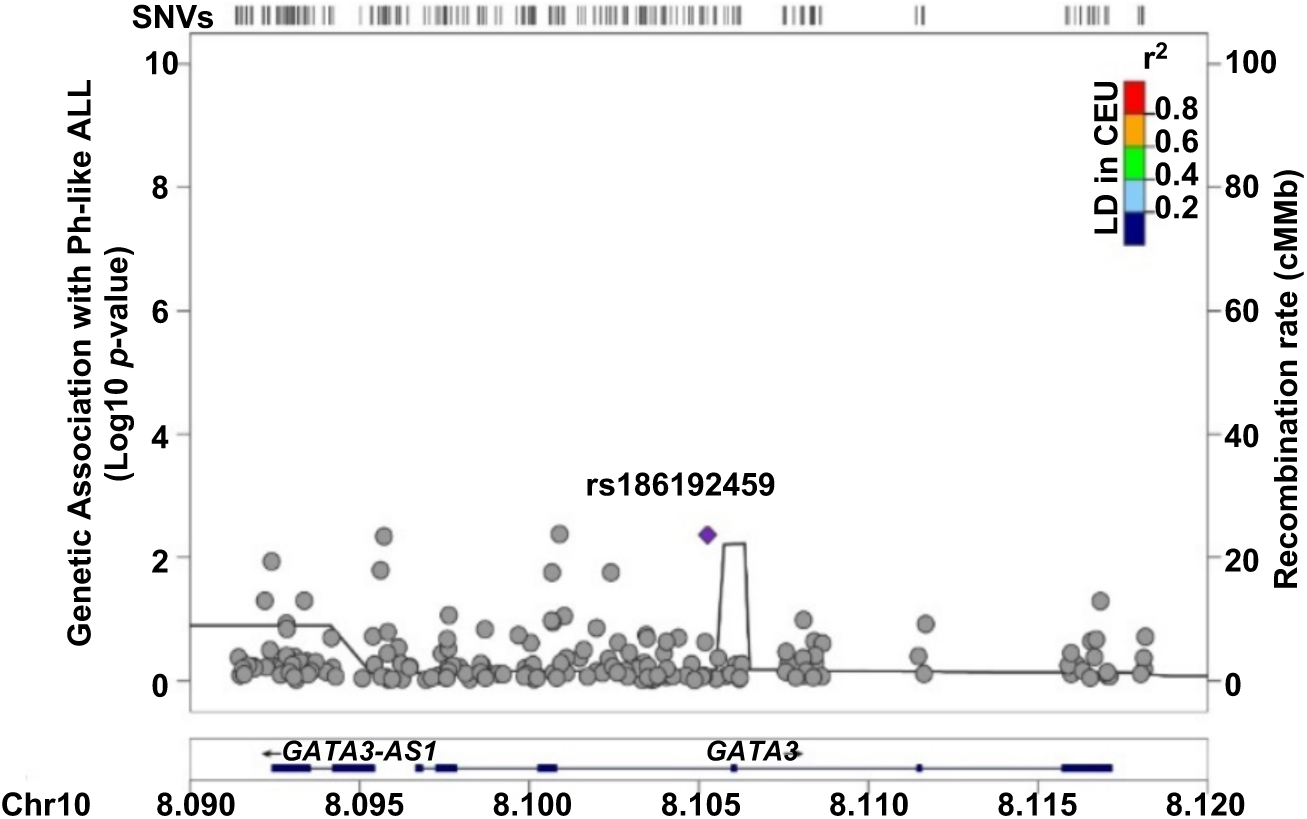
Multivariate analysis conditioning on rs3824662 revealed no independent signals reach the statistical significance to associate with Ph-like ALL susceptibility at the *GATA3* locus.

**Supplementary Figure 3.**
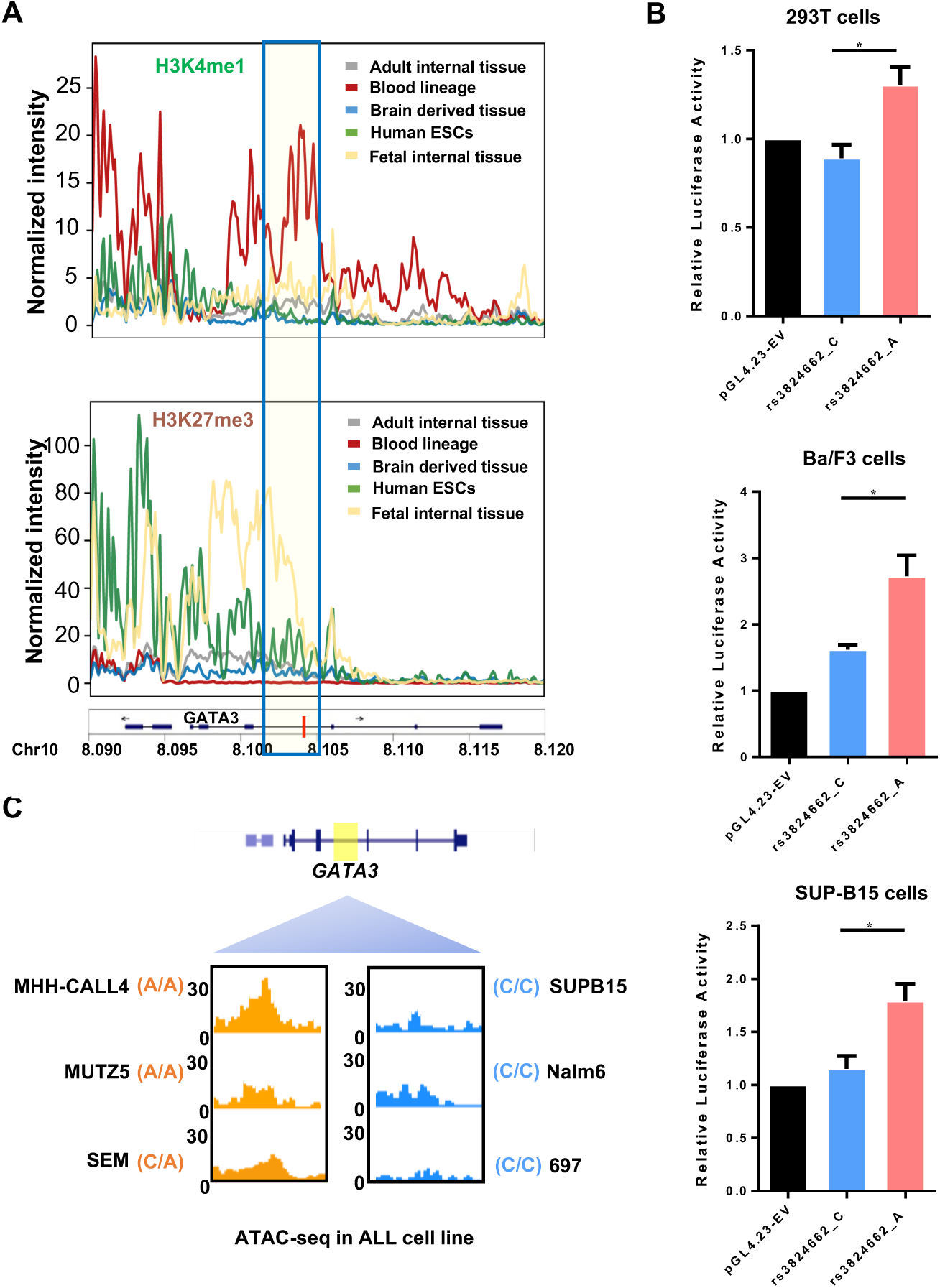
Histone signal, enhancer reporter assay and ATAC-seq analysis show rs3824662 risk A allele is associated with enhancer activity and open chromatin status in wildtype human cells, human blood tissue and human ALL cell lines. **A.** Normalized intensity of H3K4me1 and H3K27me3 signal in 42 human tissues from ROADMAP data. **B.** Luciferase reporter assay comparing the enhancer activities of the fragments containing either the rs3824662 risk A allele or wildtype C allele in human normal 293T, mouse Ba/F3 and human SUP-B15 ALL cells. T bars indicate standard deviations(student t-test: *p* value =0.0167 for 293T; *p* value =0.0138 for Ba/F3; *p* value =0.0136 for SUP-B15). **C.** Open chromatin status at the rs3824662 locus in ALL cell lines representative of different subtypes, as determined using ATAC-seq. The window represents a 2kb region flanking rs3824662. MHH-CALL4 and MUZT5 are *CRLF2*-rearranged with A/A genotype at the *GATA3* SNP; SEM is *KTM2A* rearranged and with the C/A genotype, and the other three ALL cell lines have wiltype C/C genotype (SUPB15 is BCR-ABL1 ALL, Nalm6 is *DUX*4-rearranged, and 697 is *TCF3-PBX1* ALL).

**Supplementary Figure 4.**
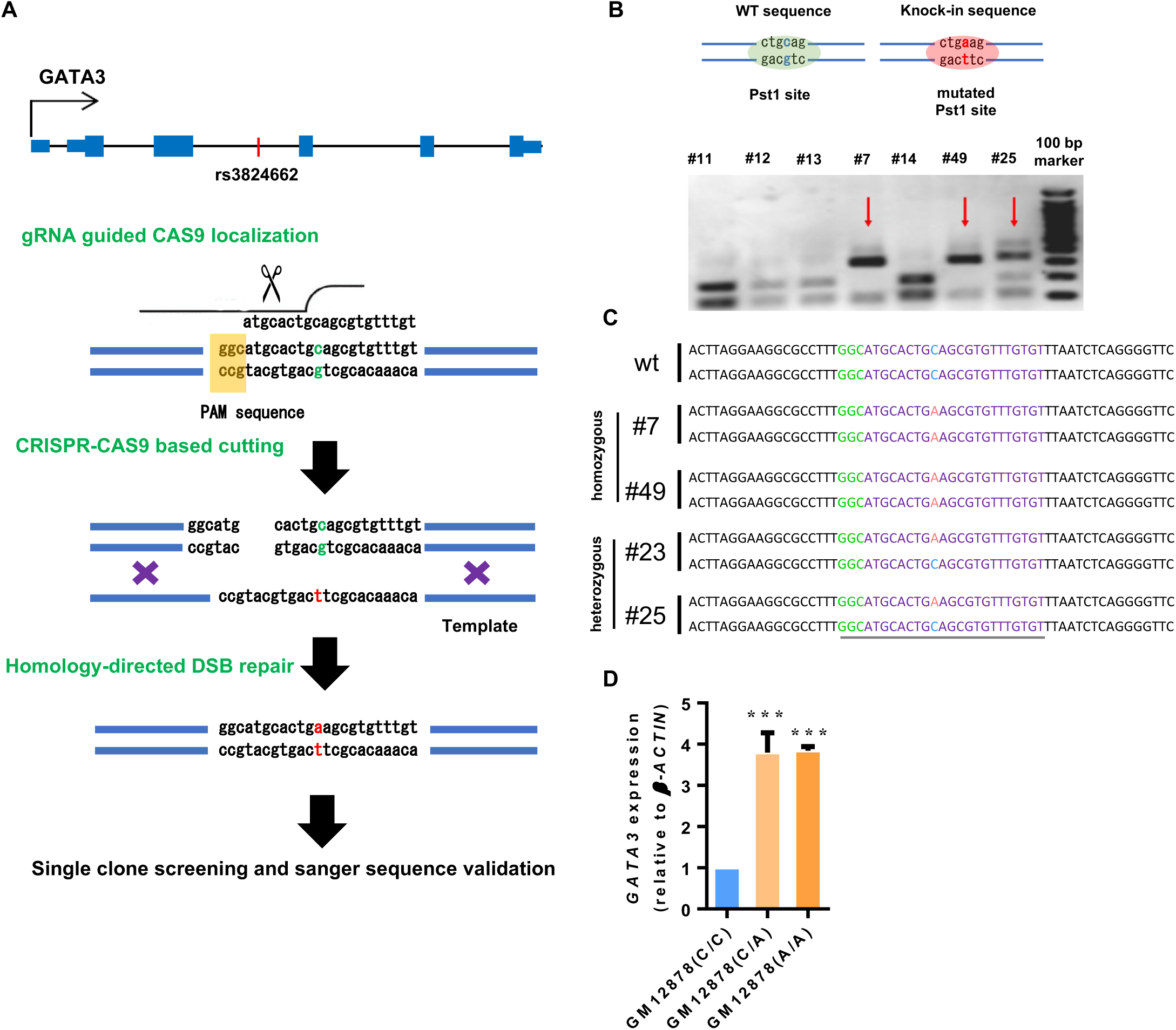
Knock-in of the rs3824662 risk A allele in GM12878 cell by using CRISPR/Cas9 editing. **A**, CRISPR design for knock-in. A 120nt template single-strand DNA containing rs3824662 A allele and flanking sequence was used as the donor for homology-directed repair with CRISPR-Cas9 induced cutting sites. **B**, Pst1 restriction enzyme is used to screen GM12878 clones with homozygous or heterozygous genotype at rs3824662. **C**, Sanger sequence results of four successful CRISPR knock-in GM12878 clones. Clone #7 and #49 had knock-in in both alleles; clone #23 and #25 had knock-in in one allele. **D**, Real time qPCR of *GATA3* expression in engineered GM12878 cells with wildtype, heterozygous, or homozygous genotype at rs3824662 (*p* value =0.0043 for C/A clones and *p* value <0.0001 for A/A clones by student *t*-test).

**Supplementary Figure 5.**
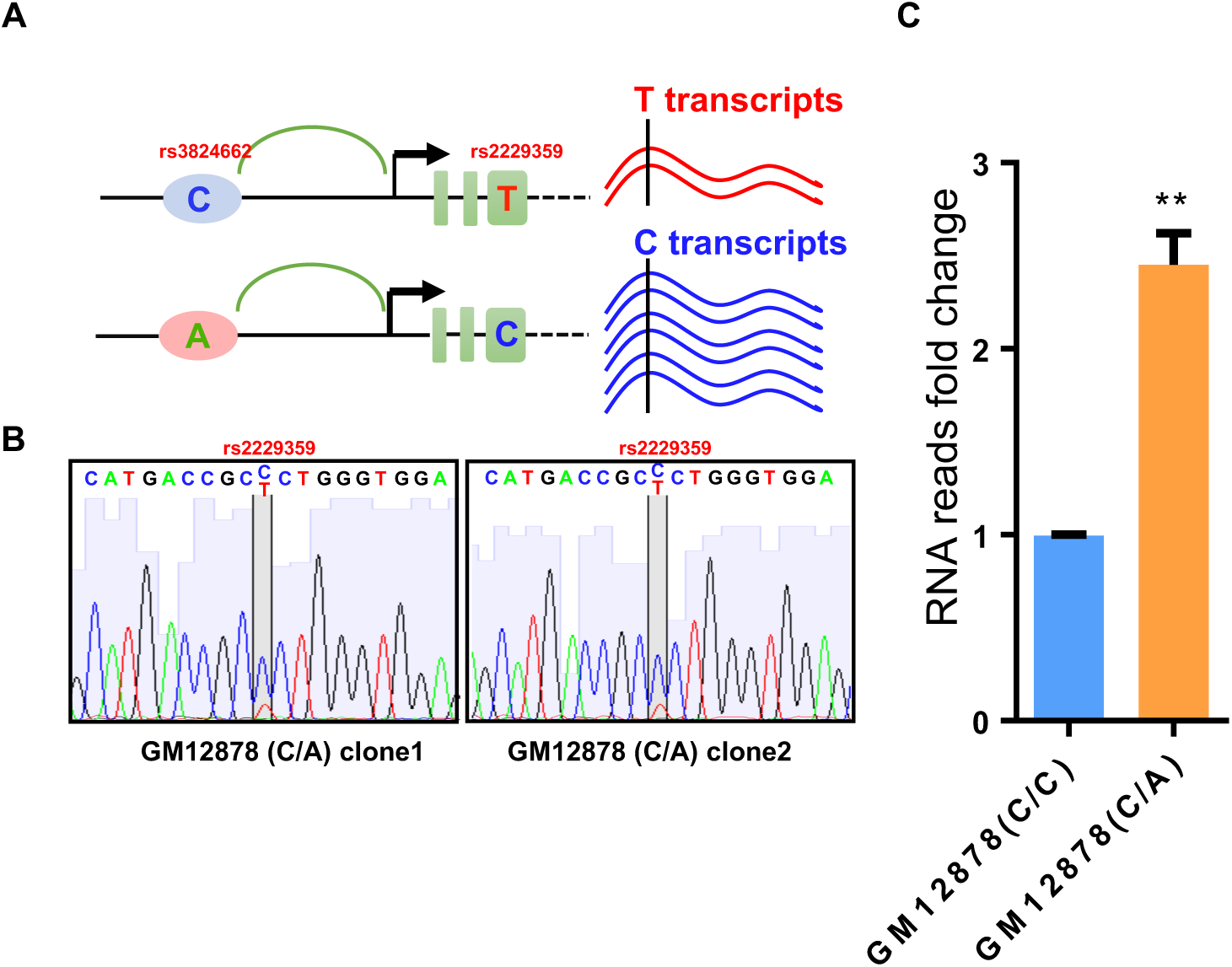
Design and detection of allelic-bias on *GATA3* gene expression in GM12878 heterozygous clones. **A.** GM12878 cells harbor a nonsynonymous variant (rs2229359 T/C) in *GATA3* 3rd exon, we performed PCR and Sanger sequencing and observed that the T allele at rs2229359 and A allele at rs3824662 are from the same allele. Therefore, allelic expression derived from rs2229359 would directly inform the differential transcription activation effects of the A *vs*. C allele at rs3824662 in engineered GM12878 clones. **B.** Sanger sequencing of PCR products of GATA3 3^rd^ exon cDNA shows allelic expression of GATA3 in two GM12878 heterozygous clone cells by rs2229359 genotyping. **C.** Shown the transcription level associated with rs3824662-A allele vs. the transcript associated with wild type C allele (*p* value =0.0066 by student t-test).

**Supplementary Figure 6.**
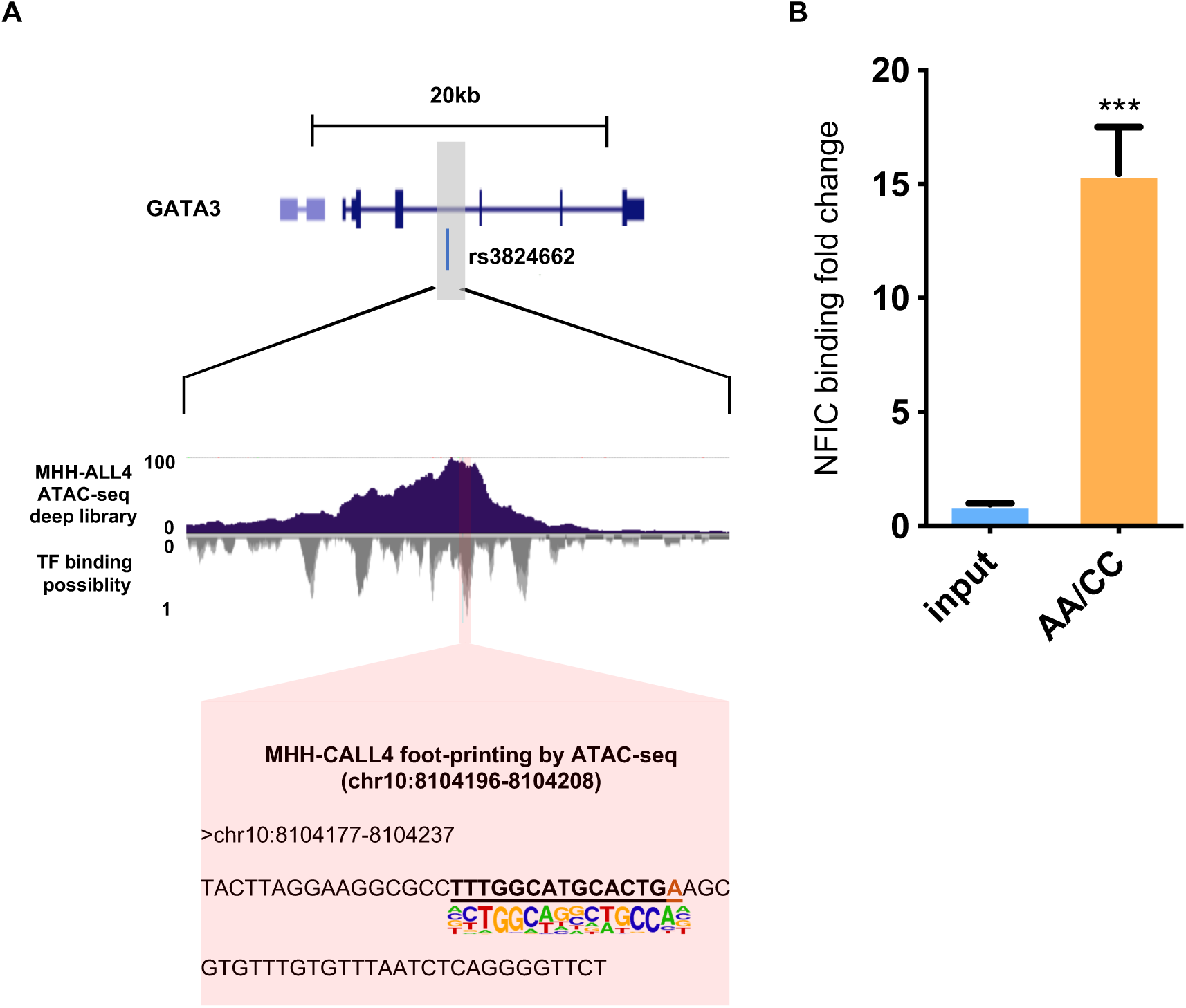
The rs3824662 risk A allele benefits NFIC binding. **A**, Footprinting analysis using high-resolution ATAC-seq data showed that the NFIC binding motif is only identified in MHH-CALL4 Ph-like ALL cell line (A/A at rs3824662), but not in GM12878 (WT) cell line. **B**, Recruitment of NFIC binding in engineered GM12878 cells with the GM12878 (A/A) compared to GM12878 (C/C) cells, measured by ChIP-qPCR (*p* value =0.0003 by student t-test).

**Supplementary Figure 7.**
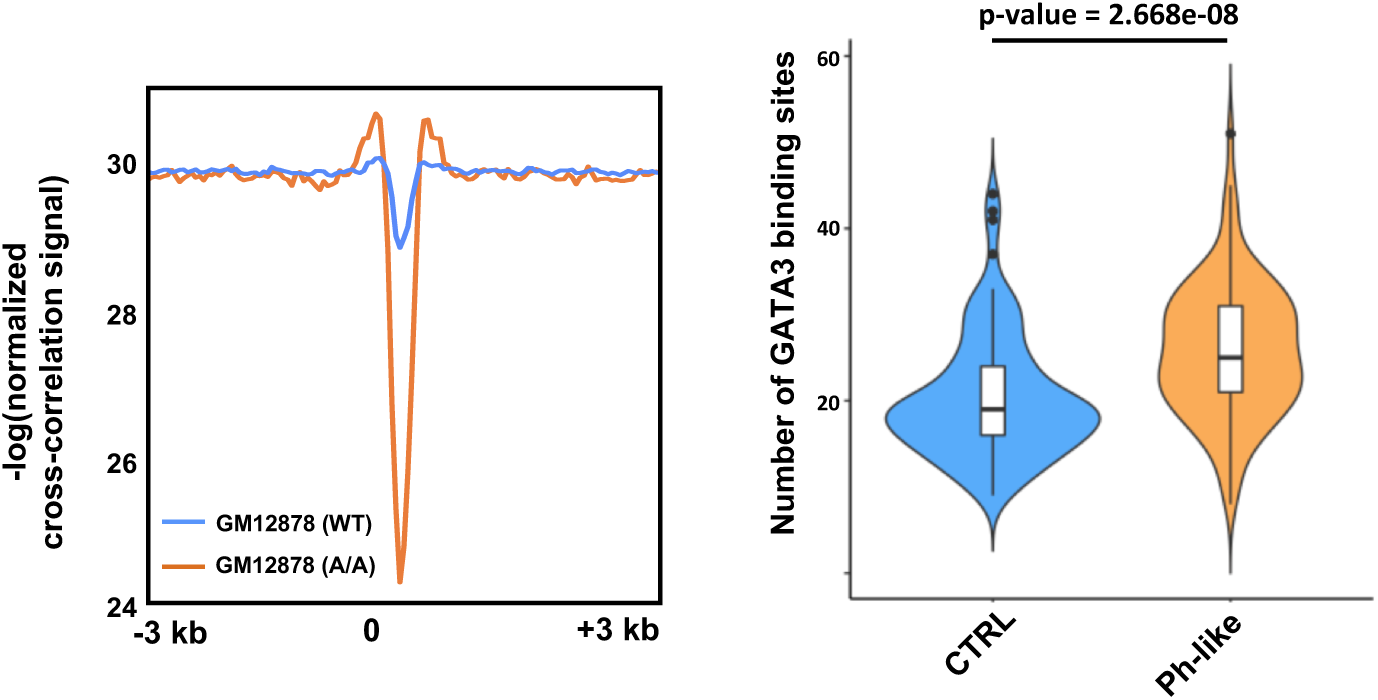
rs3824662 risk A allele induced GATA3 binding locus in the genome are devoid of nucleosomes and enriched in Ph-like genes. **A**, Nucleosome position surrounding GATA3 binding peaks in GM12878 (WT) and engineered GM12878 (A/A) cells. Y axis indicates nucleosome position probability computed from ATAC-Seq and x-axis is the 6kb window for each GATA3 binding site. **B**, Enrichment of GATA3 binding at Ph-like ALL related genes compared with random control (*p* value = 0.0003 by wilcox.test) in engineered GM12878 (A/A) cells. Ph-like genes were defined as those most differentially expressed in this subtype than other ALL, as described previously (Roberts et al 2014).

**Supplementary Figure 8.**
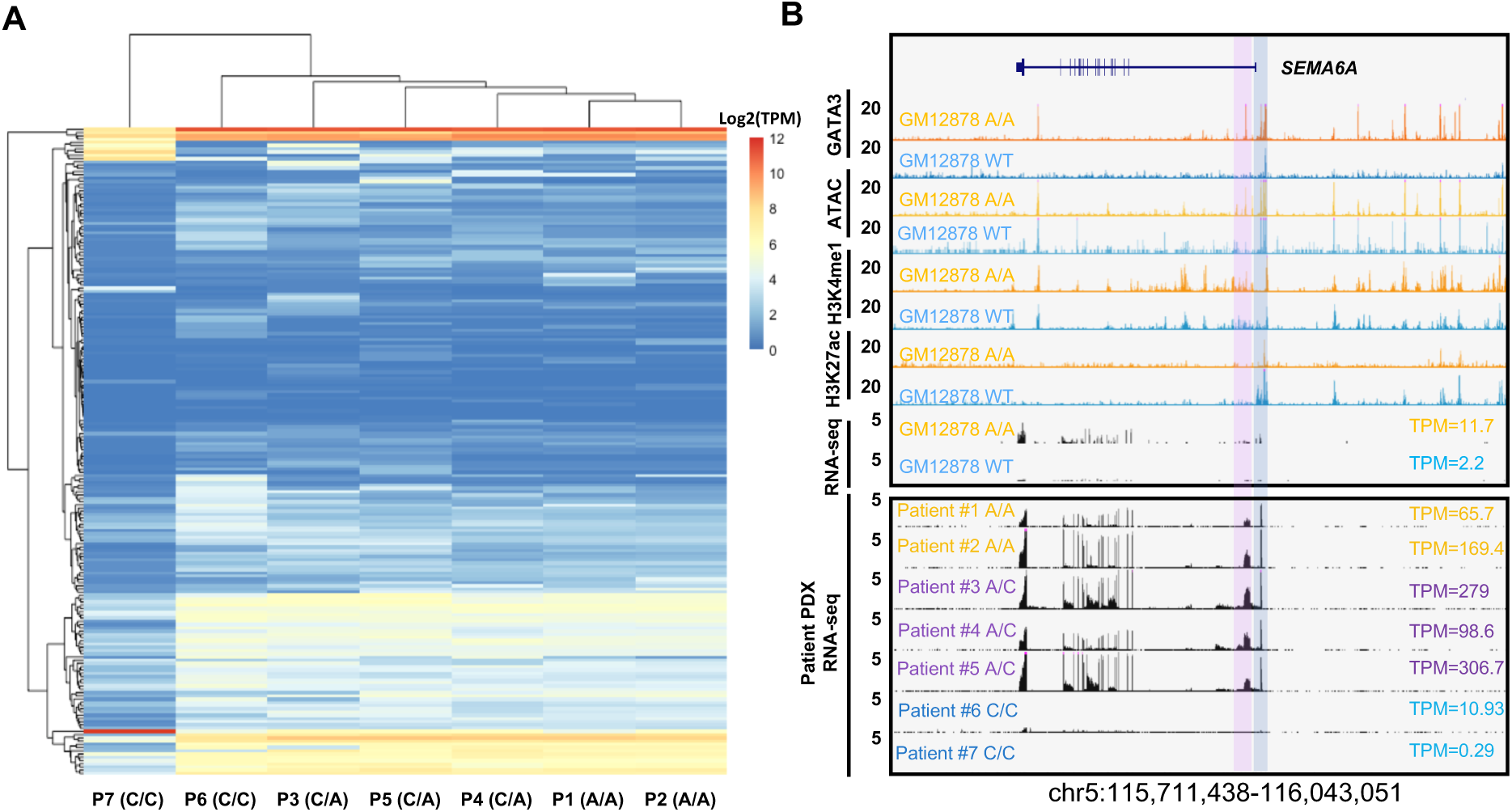
Risk A allele are also associated with ALL patient sample gene expression pattern. **A. G**lobal gene expression clustering by normalized TPM shows patient sample containing A allele are clustered together (k means = 100). **B**, Ph-like gene *SEMA6A* is highly expressed in engineered GM12878 (A/A) cells (upper panel) and also PDX samples (bottom panel) with risk A alleles. GATA3 binding is enriched in SEMA6A promoter (TSS) and enhancer (predicted by H3K27ac signal) in engineered GM12878 (A/A) cells. Blue bar and pink bar labels promoter and enhancer, respectively.

**Supplementary Figure 9.**
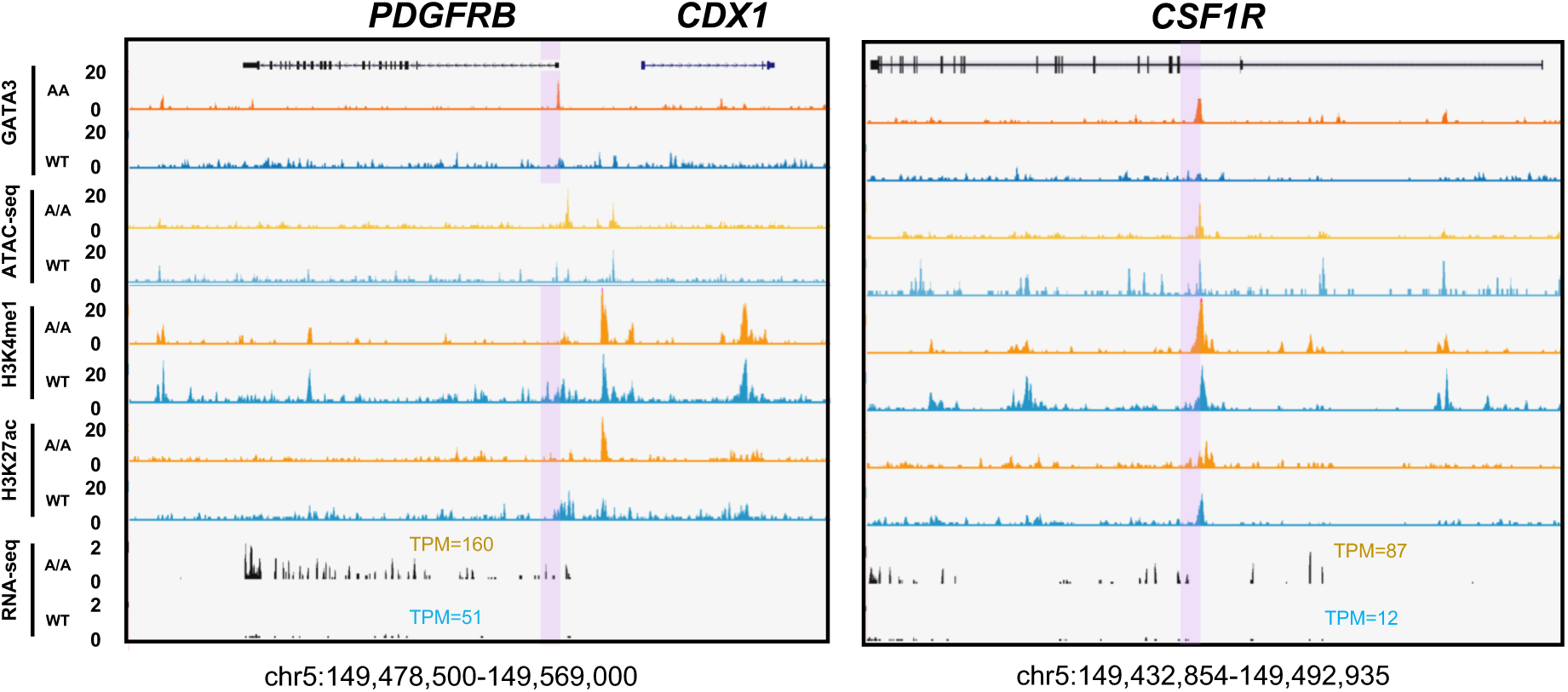
ATAC-seq, ChIP-Seq and RNA-seq for GATA3 in in GM12878 (WT) and engineered GM12878 (A/A) cells at the *PDGFRB* and *SEMA6A* loci (Read densities (y axis) normalized by sequencing depths). Pink bar indicates GATA3 binding in enhancer area.

**Supplementary Figure 10.**
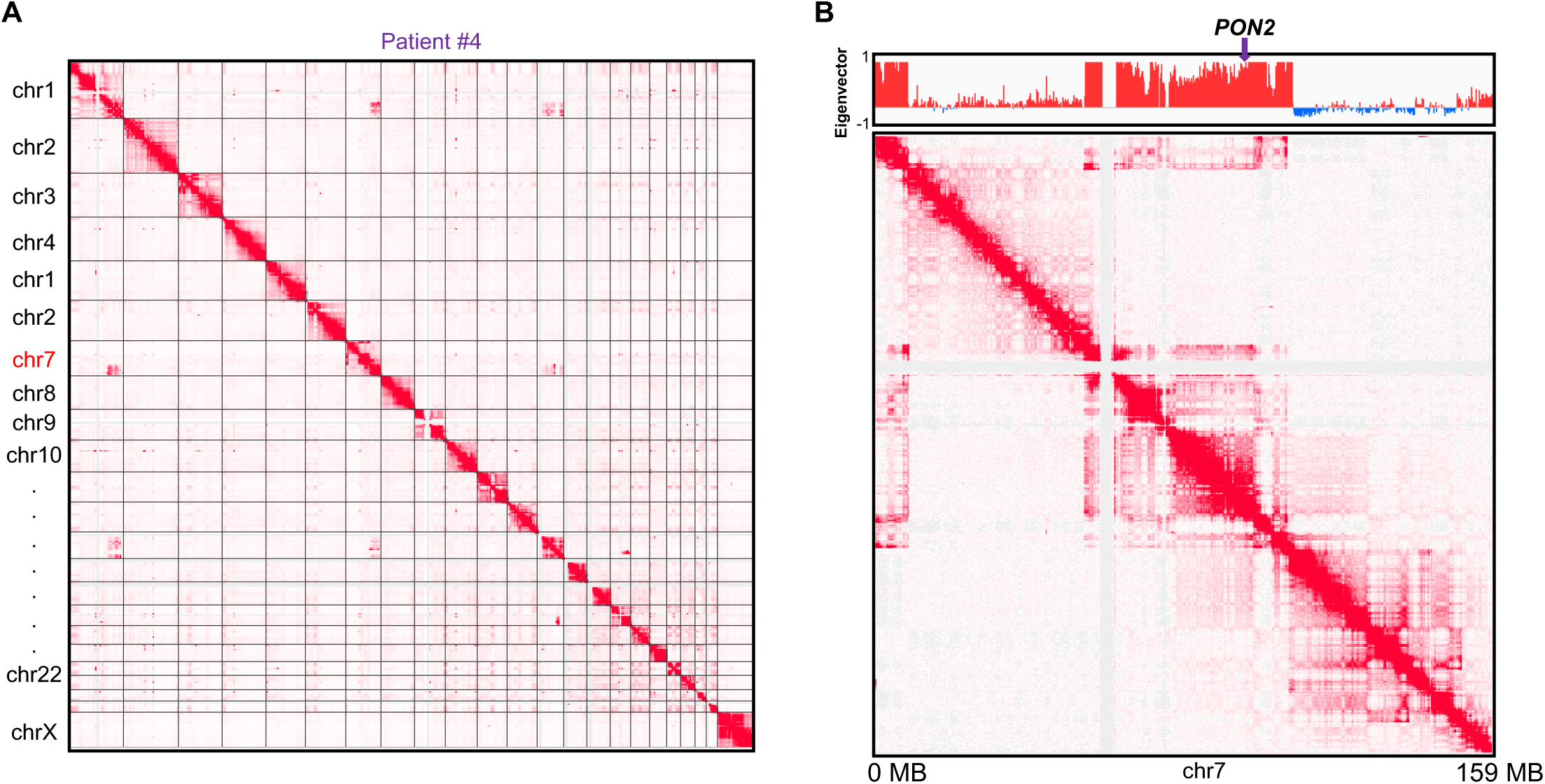
interchromosomal (**A**) and intrachromosomal (**B**) rearrangements in chromosome Patient #4 indicated by Hi-C heatmap. B up panel showed the abnormal compartment state in chr7.

**Supplementary Figure 11.**
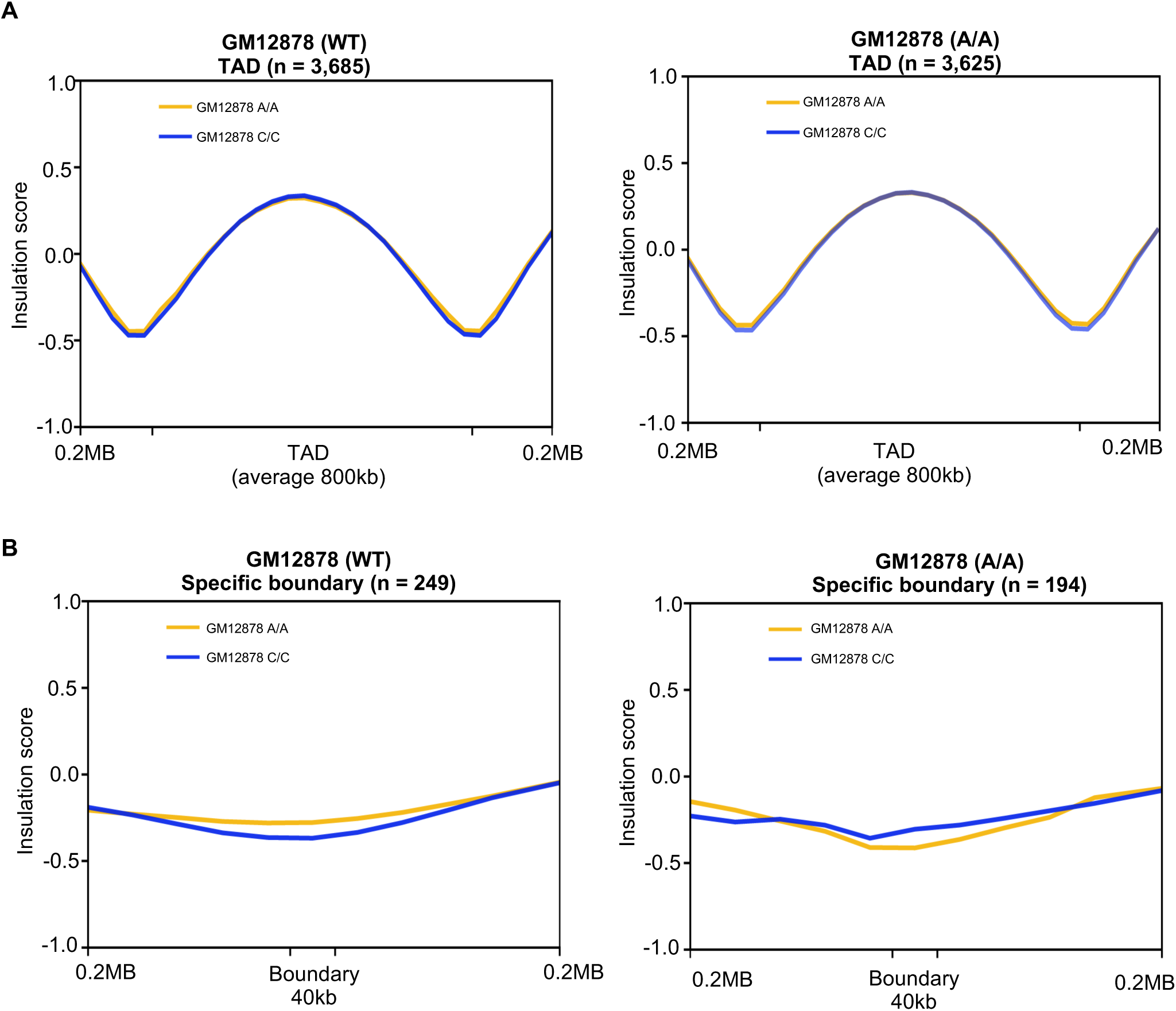
TAD structure is consistent in GM12878 (WT) and engineered GM12878 (A/A) cells. **A**, Average insulation score shows no significant difference in GM12878 cells with different rs3824662 genotype. Left panel: Insulation score from GM12878 (WT) Hi-C result (blue line) and engineered GM12878 (A/A) Hi-C result (yellow line) in GM12878 (WT) TADs. Right panel: Insulation score from GM12878 (WT) Hi-C result (blue line) and engineered GM12878 (A/A) Hi-C result (yellow line) in GM12878 (A/A) TADs. **B**, Average insulation score shows no significant difference in GM12878 (WT) Hi-C (blue line) and engineered GM12878 (A/A) Hi-C (yellow line) in GM12878 (WT) specific TAD boundaries. Left panel: Insulation score from GM12878 (WT) Hi-C (blue line) and GM12878 (A/A) Hi-C (yellow line) in GM12878 (C/C) TAD boundaries. Right panel: Insulation score from GM12878 (WT) Hi-C (blue line) and GM12878 (A/A) Hi-C (yellow line) in GM12878 (WT) TAD boundaries.

**Supplementary Figure 12.**
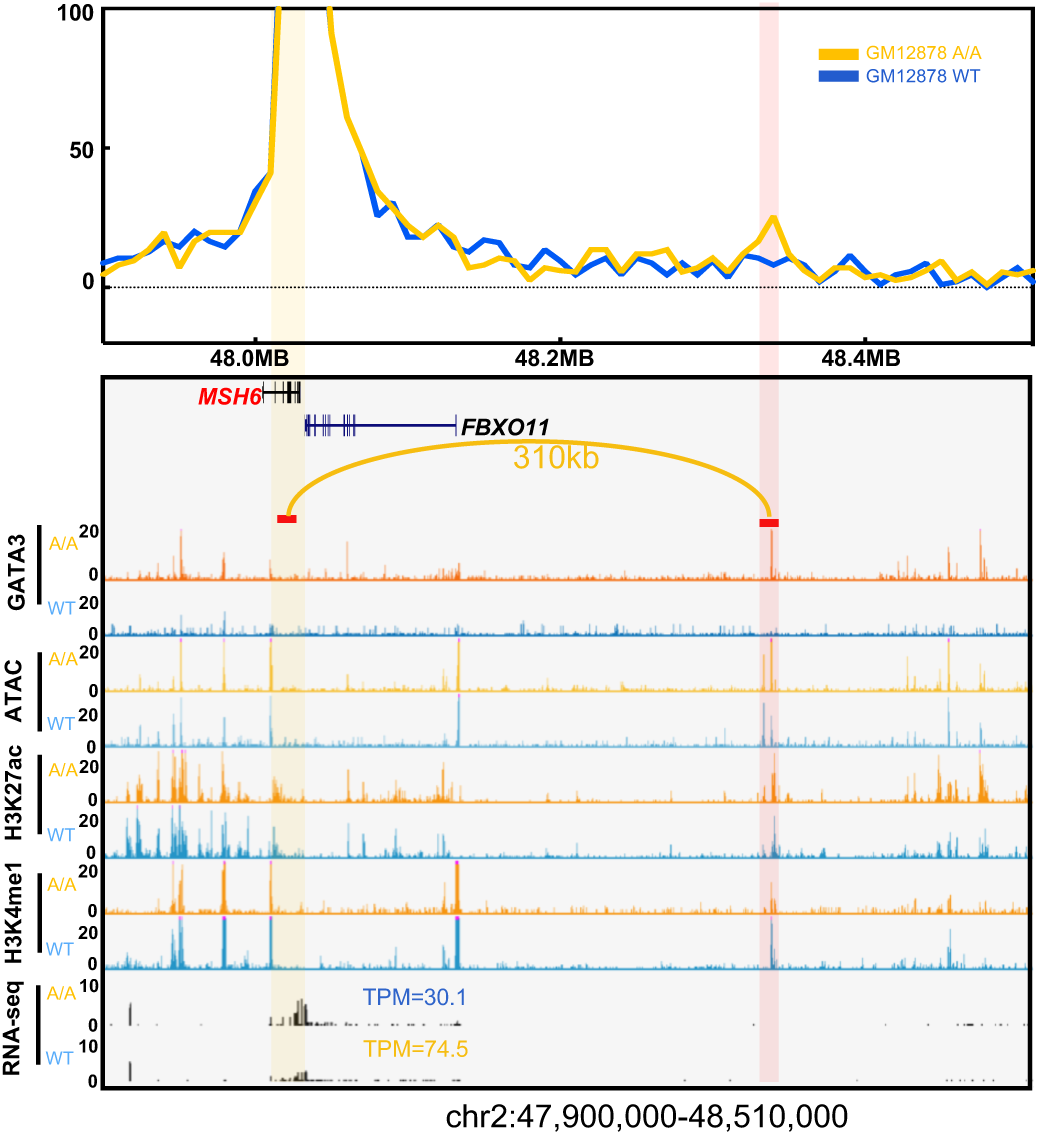
Virtual 4-C analysis in 10kb resolution shows there is a A/A genotype-specific chromatin looping between the *MSH6* locus (yellow bar) and one predicted enhancer 310kb away (pink bar) in engineered GM12878 (A/A) cells (related to figure 4a).

**Supplementary Figure 13.**
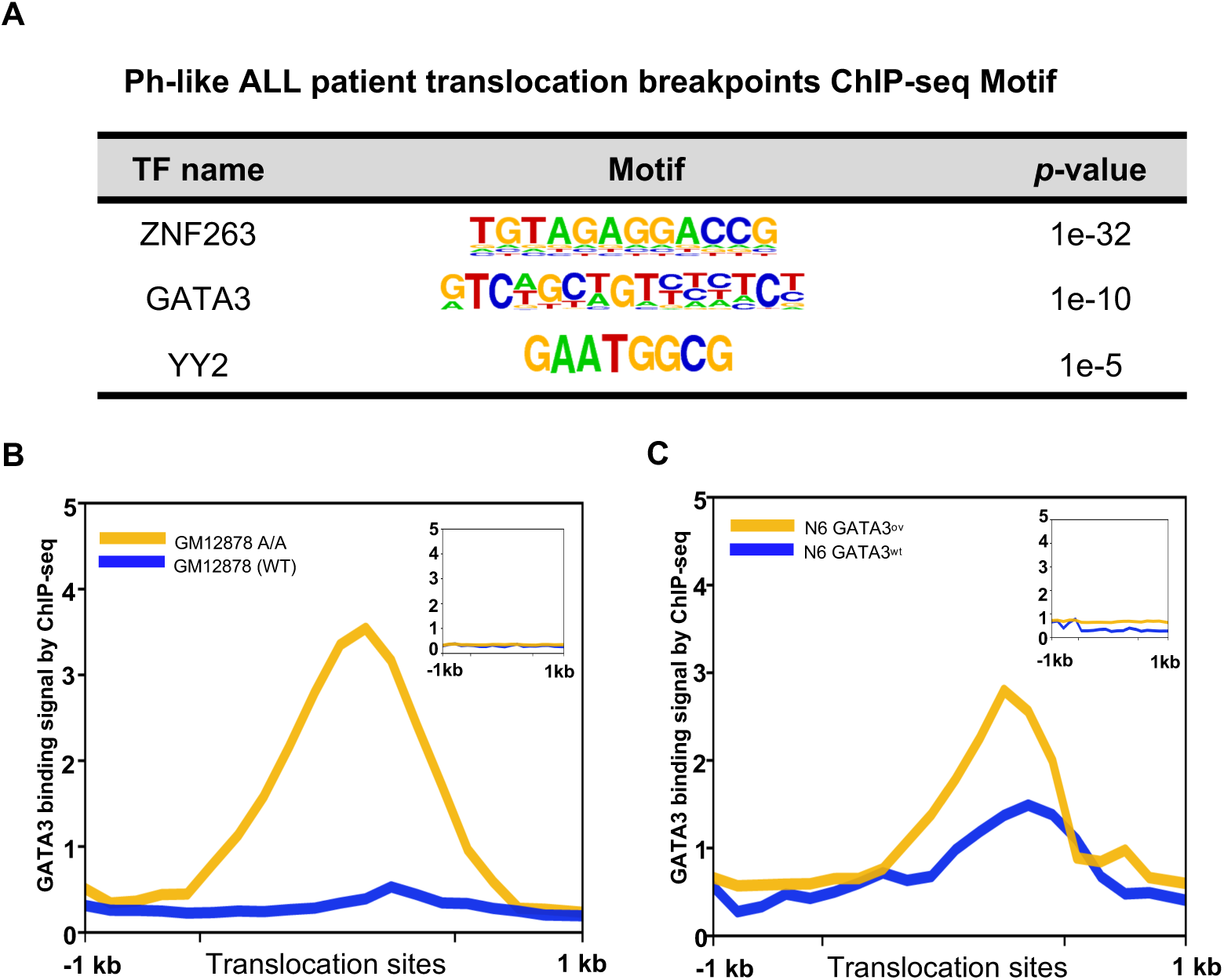
GATA3 involved in translocation in Ph-like ALL. **A**, Motif enrichment analysis of Ph-like ALL patient translocation breakpoints genomic regions. **B**, GATA3 binding signal (200bp bin) in Ph-like ALL patient translocation breakpoints region by ChIP-seq (+/- 1kb) in engineered GM12878 (A/A) (yellow) and GM12878 (WT) cells (blue). Upright panel is GATA3 binding signal in 1000 random genomic regions in GM12878 A/A and WT cells. **C**, GATA3 binding signal (200bp bin) in Ph-like ALL patient translocation breakpoints region by ChIP-seq (+/- 1kb) in Nalm6 GATA3^ov^ (yellow) and Nalm6 GATA3^wt^ (blue) cells. Upright panel is GATA3 binding signal in 1000 random genomic regions in Nalm6 GATA3^ov^ (yellow) and Nalm6 GATA3^wt^ (blue) cells.

**Supplementary Figure 14.**
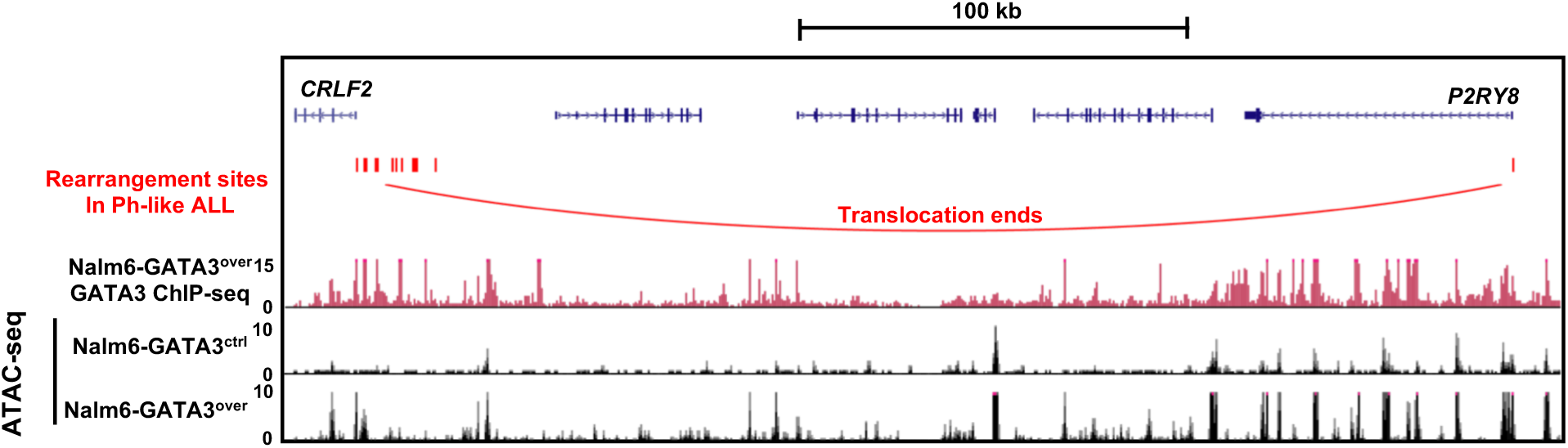
GATA3 ChIP-seq and ATAC-seq in Nalm6 with or without ectopic *GATA3* expressed at the *CRLF2* locus. Red vertical bars indicate the rearrangement hotspots in *CRLF2*-positive Ph-like ALL. ChIP-seq and ATAC signal intensities were normalized according to their sequencing depths.

**Supplementary Figure 15.**
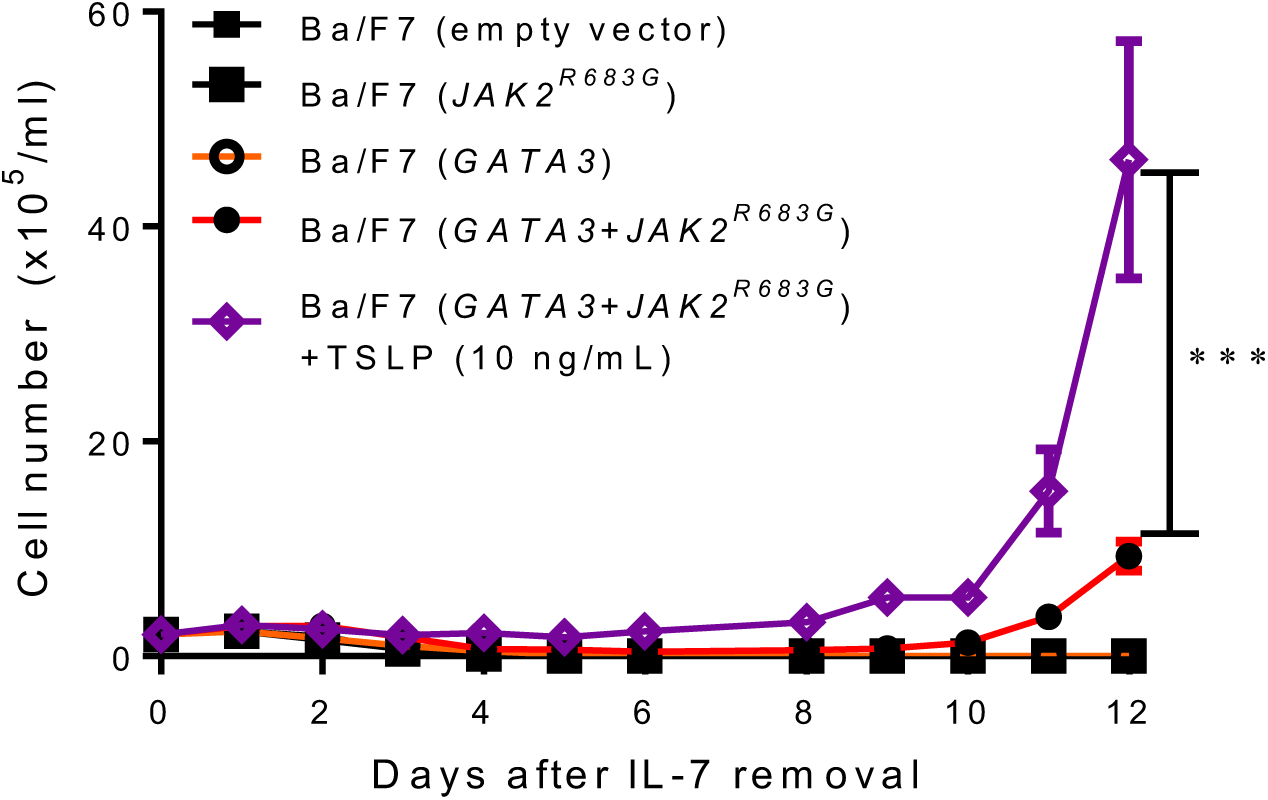
IL3-independent growth of Ba/F7 cells transduced with *GATA3* alone, *JAK2^R683G^* alone, *GATA3* with *JAK2^R683G^*, or empty vector control. All the experiments were performed in triplicates (*p* value < 0.001 by 2way ANOVA). Ba/F7 cells with *GATA3* and *JAK2^R683G^* were treated with or without 10 ng/ml TSLP. All the experiments were performed in triplicate for three times independently (*p* value < 0.001 by 2way ANOVA).

**Supplementary Figure 16.**
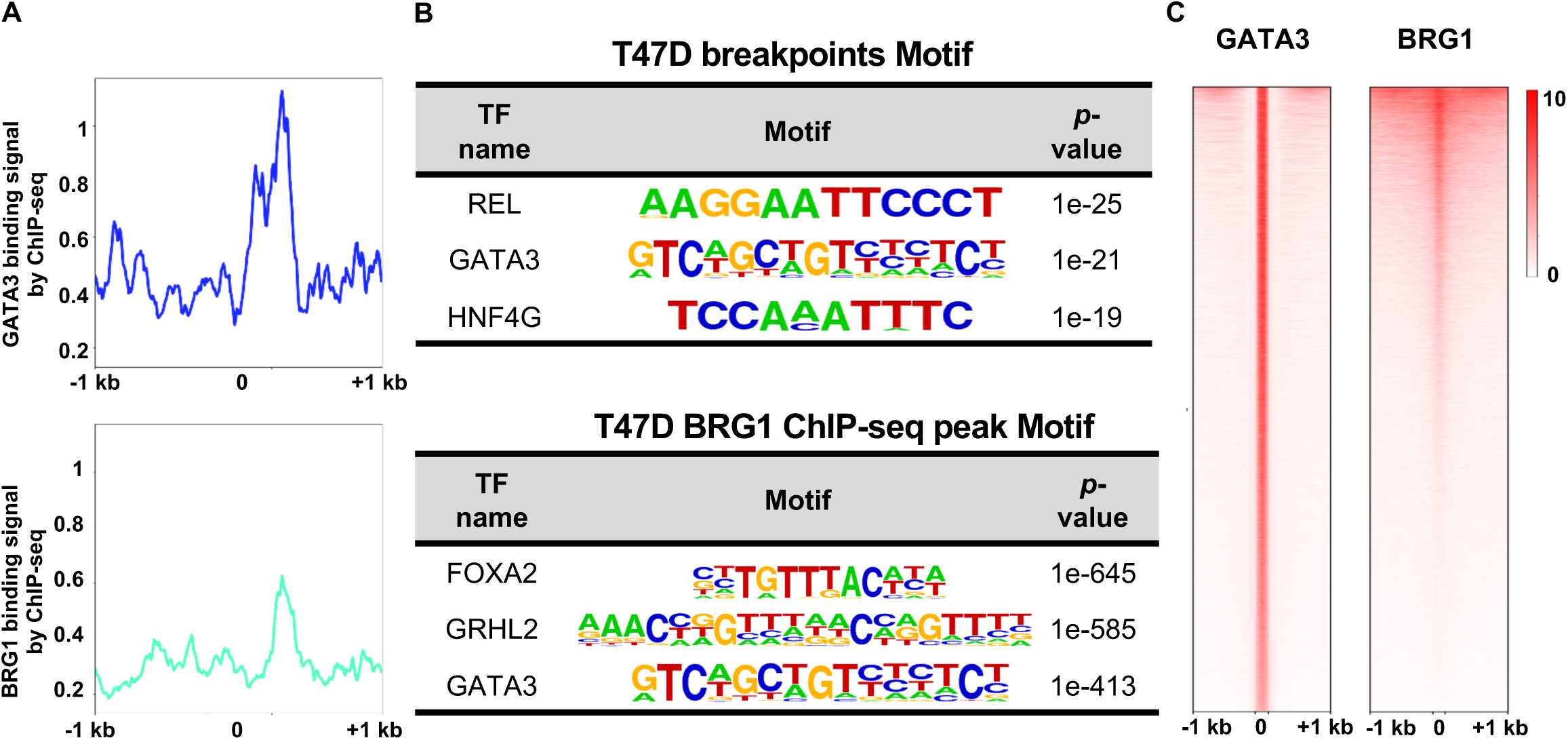
GATA3 involved in translocation in breast cancer cell line. **A**, GATA3 and BRG1 binding signal in T47 breakpoints region by ChIP-seq (+/- 1kb). **B**, Motif enrichment analysis of T47D breakpoints genomic regions and BRG1 ChIP-seq peak region. **C**, Heatmap of GATA3 and BRG1 binding signal in GATA3 binding peaks (+/- 1 kb) in T47D cell.

